# The lupus susceptibility locus Sgp3 encodes the suppressor of endogenous retrovirus expression SNERV

**DOI:** 10.1101/487231

**Authors:** Rebecca S. Treger, Scott D. Pope, Yong Kong, Maria Tokuyama, Manabu Taura, Akiko Iwasaki

## Abstract

Elevated endogenous retrovirus (ERV) transcription and anti-ERV antibody reactivity are implicated in lupus pathogenesis. Overproduction of non-ecotropic ERV (NEERV) envelope glycoprotein, gp70, and resultant nephritis occur in lupus-prone mice. However, a NEERV repressor has not been identified to test if this association is causal. Here we identified suppressor of <u>N</u>E<u>ERV</u> *(Snerv)* 1 & 2, Krüppel-associated box zinc finger proteins (KRAB-ZFP) that repressed NEERV by binding the NEERV long terminal repeat to recruit the transcriptional regulator KAP1. Germline *Snerv1/2* deletion increased activating chromatin modifications, transcription, and gp70 expression from NEERV loci. F1 crosses of lupus-prone NZB and 129 mice to *Snerv1/2^−/−^* mice failed to restore NEERV repression, demonstrating that loss of SNERV underlies the lupus autoantigen gp70 overproduction that promotes nephritis in susceptible mice. Increased ERV expression in lupus patients was inversely correlated with expression of three putative ERV-suppressing KRAB-ZFP, suggesting that KRAB-ZFP-mediated ERV misexpression may contribute to human lupus pathogenesis.

## Introduction

Retroelements (RE) are mobile DNA species that compose ~40% of murine and human genomes (Lander et al., 2001; Waterston et al., 2002). Although generally silenced, these elements can cause insertional mutagenesis and have diverse effects upon gene expression (Goodier, 2016). The ability to limit RE movement in the genome is fundamentally important, as transposon-mediated disruption or dysregulation of genes contributes to more than 100 human diseases, including hemophilia and leukemia (Goodier, 2016; Hancks and Kazazian, 2016; Kazazian and Moran, 2017). Endogenous retroviruses (ERV) are RE formed by the remnants of past retroviral infection that have accumulated in the genome over millennia. Many ERV retain transposition potential and are responsible for ~10% of spontaneous mutations in inbred mice (Kazazian and Moran, 1998; Maksakova et al., 2006). More recently acquired ERV have retained envelope-coding regions, in addition to structural genes that encode the gag matrix, protease, and polymerase (Kozak, 2014). These proviral ERV are located throughout the genomes of inbred mouse strains (Coffin et al., 1989).

As with exogenous retroviruses, infectious ERV, originally identified in constitutively viremic mouse strains, are appreciated for their role in malignant transformation (Kassiotis, 2014; Kozak, 2014). Additionally, in certain immune deficient murine backgrounds and cancer cell lines, ERV transcripts from mouse-tropic (i.e. ecotropic) and non-ecotropic ERV (NEERV) loci recombine to generate infectious ERV (Ottina et al., 2018; Young et al., 2012; Yu et al., 2012). Thus, transcriptional silencing of genomic ERV sequences is a critical layer of defense from active retrotransposition, restoration of infectivity, and insertional mutagenesis leading to oncogenesis.

RE loci are targeted by epigenetic modifications that result in establishment and maintenance of transcriptional repression (Macfarlan et al., 2011; Matsui et al., 2010; Rowe et al., 2013b; Wolf and Goff, 2007). This transcriptional silencing is generally initiated by Krüppel-associated box domain zinc finger proteins (KRAB-ZFP), a large family of DNA-binding transcriptional regulators in vertebrates (Ecco et al., 2017). KRAB-ZFP can recognize and bind to DNA sequences common in RE families through their C-terminal zinc fingers and recruit KRAB-associated protein-1 (KAP1) through the N-terminal KRAB domain to form a scaffold around which transcriptional silencing machinery can assemble (Ecco et al., 2017; Rowe et al., 2013a; Rowe et al., 2010). ZFP809 binds to and silences ecotropic ERV loci in this manner (Wolf and Goff, 2009; Wolf et al., 2015). However, a specific KRAB-ZFP repressor responsible for silencing NEERV transcripts in mice has not yet been identified.

While under much speculation, the role of ERV dysregulation in the pathogenesis of autoimmune disease is not well established. Elevated transcription of human ERV (HERV) loci and antibody reactivity to HERV proteins occurs in many autoimmune diseases (Grandi and Tramontano, 2018; Gröger and Cynis, 2018). In systemic lupus erythematosus (SLE) patients, hypomethylation of HERV loci and antibody reactivity to HERV and retroviral (HIV-1, HTLV-1) proteins are implicated in SLE pathogenesis (Blomberg et al., 1994; Hishikawa et al., 1997; Mellors and Mellors, 1976; Nakkuntod et al., 2013; Perl et al., 1995; Wu et al., 2015). This association between HERV dysregulation and SLE pathogenesis is further strengthened by murine models of spontaneous lupus, where NEERV envelope glycoprotein gp70 is a major autoantigen promoting lupus nephritis (Baudino et al., 2008; Ito et al., 2013; Yoshiki et al., 1974). Yet the association between HERV dysregulation and SLE remains tentative: HERV are poorly annotated in the genome and knowledge about HERV transcriptomes is limited; specific factors that modulate HERV expression in SLE patients have not been identified; and molecular mechanisms linking HERV dysregulation to SLE pathogenesis have not been defined (Nelson et al., 2014). Even in murine lupus models, the gene and mechanism responsible for NEERV dysregulation is not known. The Gross virus antigen 1 *(Gv1)* locus in 129 strains and the serum gp70 production 3 *(Sgp3)* locus in lupus-prone New Zealand Black (NZB) and New Zealand White (NZW) strains both drive elevated NEERV expression, a major hallmark of disease (Andrews, 1978; Baudino et al., 2008; Izui, 1979). While the *Sgp3* and *Gv1* loci have been mapped by QTL analyses to an interval on chr13 (Laporte et al., 2003; Oliver and Stoye, 1999), the identity of the gene(s) responsible for the gp70 overexpression remain unknown.

In this study, we identified the KRAB-ZFP genes within the *Sgp3* and *Gv1* loci that are responsible for silencing of NEERV transcripts. We also examined HERV mRNA expression in the peripheral blood mononuclear cells (PBMC) of SLE patients and found putative HERV-suppressing KRAB-ZFP genes whose expression inversely correlated to that of HERV. Our findings suggest that a similar defect in HERV repression may promote human lupus pathogenesis.

## Results

### NEERV transcription is globally increased in C57BL/6N, but not C57BL/6J, lymphocytes and bone marrow-derived macrophages

In experiments to test innate viral sensors involved in control of ERV, we found that steady-state lymphocyte NEERV envelope mRNA and protein expression from xenotropic (Xmv) and polytropic (Pmv) loci differed by background substrains: NEERV expression was increased in C57BL/6N (B6N) compared to C57BL/6J (B6J) (Figure 1A-B). These substrains were separated only ~70 years ago, and a number of SNPs differentiate these substrains (Mekada et al., 2008; Simon et al., 2013). To identify B6N and B6J transcriptome differences, RNA-sequencing was carried out in naïve CD4^+^ T cells. To map sequencing reads to unique proviral ERV loci, we developed an analysis pipeline in which we used a list of proviral ERV loci obtained from Jern et al. (Jern et al., 2007) in combination with an algorithm composed of stringent filtering criteria adapted from Schmitt et al. (Schmitt et al., 2013). The major transcriptional difference between B6N and B6J mice was a global increase in B6N Xmv, Pmv, and modified polytropic (Mpmv) NEERV transcripts, with minimal impact on the ecotropic ERV, *Emv2,* or on other cellular genes (Figure 1C). NEERV transcription was increased regardless of whether reads were mapped to unique NEERV loci (Figure 1C) or to NEERV long terminal repeat (LTR) families (Figure 1D). This phenotype was penetrant across various cell types, including lymphocytes, total bone marrow, bone marrow-derived macrophages (BMDM), and embryonic stem cells (Figure S1A-B). Indeed, all uniquely mappable NEERV loci and NEERV LTR families were also increased in RNA-sequencing of B6N BMDM, compared to B6J BMDM (Figure 1E-F). Across cell types, increased B6N NEERV expression was also highly specific to this RE family; by mapping to unique loci or entire repeat families, the expression of long interspersed nuclear elements 1 (LINE1) and other LTR family by RT-qPCR or RNA-sequencing was unchanged in either naïve CD4 T cells or BMDM (Figure 1C,E & Figure S1C-D). Thus, B6N mice expressed elevated levels of NEERV mRNA and envelope protein compared to B6J mice.

**Figure 1.**
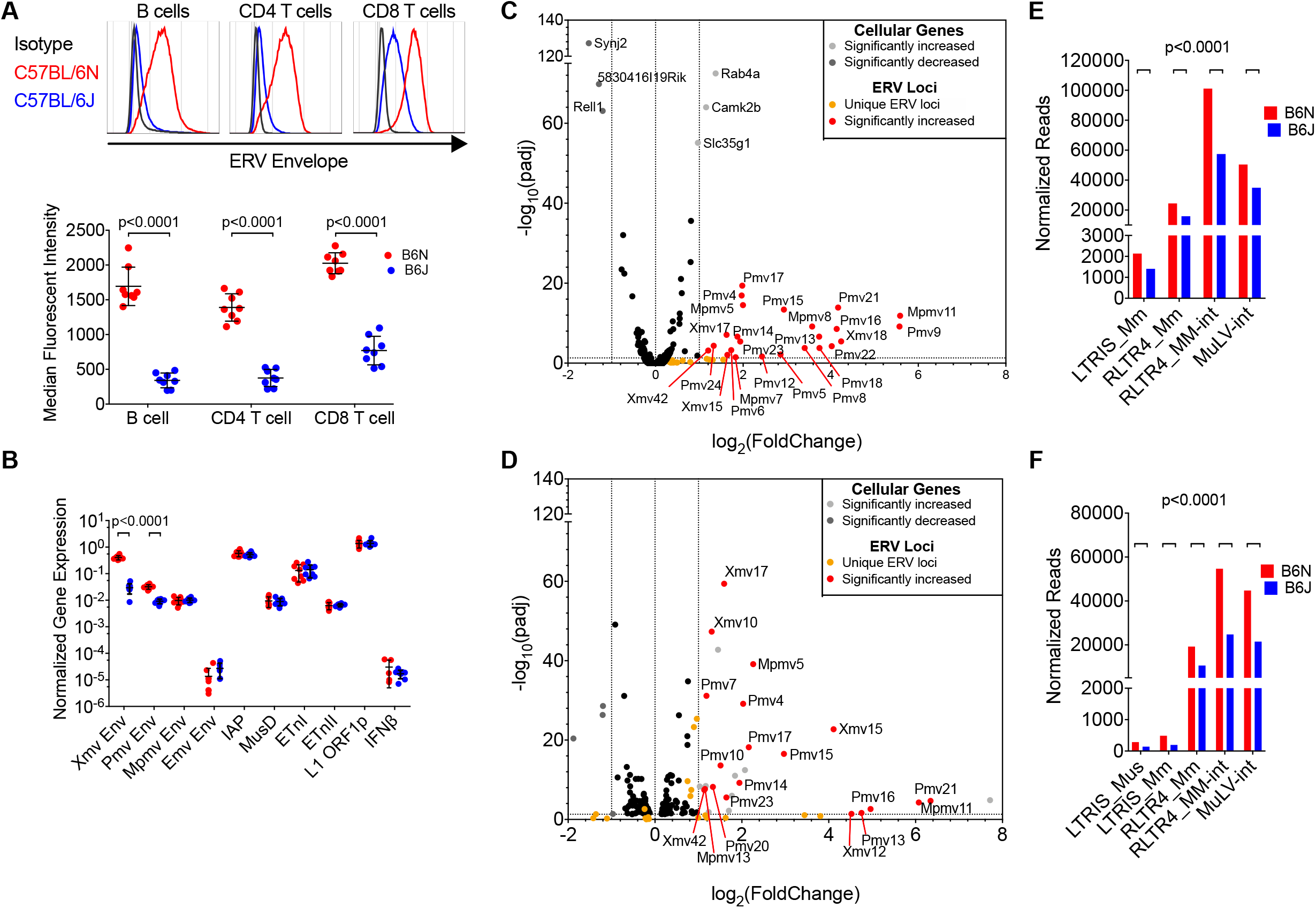
NEERV transcription is globally increased in C57BL/6N, but not C57BL/6J, lymphocytes and bone marrow-derived macrophages. **(A)** Representative histogram and calculated MFI of ERV envelope protein expression detected via FACS on the surface of peripheral blood B cells, CD4^+^ T, and CD8^+^ T lymphocytes from adult C57BL/6N (B6N) and C57BL/6J (B6J) mice. Each histogram or point represents an individual mouse and mean and standard deviation are plotted. **(B)** RT-qPCR of RNA from total splenocytes from B6N (n=8) and B6J (n=8) mice. Primers amplify respective envelope regions of all Xmv, Pmv, Mpmv, and Emv transcripts, the gag or polymerase regions of IAP, MusD, and ETn elements (Maksakova et al., 2009), or LINE1 ORFp1. Values were normalized to GAPDH expression. Mean and standard deviation are plotted. **(C)** Volcano plot of differentially expressed cellular genes & all 47 uniquely mappable ERV loci from mRNA sequencing of B6N and B6J naïve CD4^+^ T cells. (D) Normalized read counts mapping to NEERV LTR families using the RepEnrich alignment strategy from mRNA sequencing of naïve CD4^+^ T cells. (E) Volcano plot of differentially expressed cellular genes & all 47 uniquely mappable ERV loci from mRNA sequencing of B6N and B6J bone marrow-derived macrophages (F) Normalized read counts mapping to NEERV LTR families using the RepEnrich alignment strategy from mRNA sequencing of bone marrow-derived macrophages. Adjusted p-values in Figure 1 and Figure S1 were calculated for multiple t-tests (two-tailed) comparing B6N to B6J for each gene, corrected for the 25 independent hypotheses tested in Figure 1 and Figure S1 using the Holm-Sidák method with an alpha value of 0.05 for the entire family of comparisons. Adjusted p-values in Figure 1D & Figure 1E were calculated using DESeq2. See also Figure S1.

### Intergenic NEERV loci are enriched for activating histone modifications and depleted of repressive histone modifications in B6N bone marrow-derived macrophages

While actively transcribed regions are enriched for histone modifications including histone 3 lysine 4 trimethylation (H3K4me3) and H3K27 acetylation (H3K27Ac), RE are generally enriched for the repressive histone mark H3K9me3 (Groh and Schotta, 2017). To investigate if epigenetic silencing is perturbed at B6N NEERV loci, we performed chromatin immunoprecipitation and sequencing (ChIP-seq) of B6N and B6J BMDM for active and repressive histone marks. We mapped ChIP-seq reads to unique NEERV loci (Jern et al., 2007), including flanking upstream and downstream genomic sequences. To avoid confounding regulation of NEERV elements with regulation of genes within which they reside, we excluded 19 non-intergenic NEERV loci from analysis. We additionally mapped the ChIP-seq reads to all 71 intergenic loci that encode unique full-length viral-like 30 (VL30) elements (Markopoulos et al., 2016). VL30s are retrovirus-like LTR RE that contain gag matrix and integrase/polymerase coding regions but lack intact open reading frames. While they share many of the same structural elements as NEERV and are actively transcribed, VL30 mRNA were not differentially expressed between B6N and B6J mice (Figure S1C-D). Unlike B6N VL30 loci, intergenic B6N NEERV loci were significantly enriched for H3K4me3 and H3K27Ac (Figure 2A-B), whether mapping reads to the full-length (Figure 2A, top row) or to the first 2kb (Figure 2A, middle row) of the RE loci. Concordantly, B6N NEERV, and not VL30, loci were significantly depleted of H3K9me3, regardless of whether reads were mapped to the full-length or to the first 2kb of the RE loci. There were no differences in activating and repressive marks in the region immediately upstream (1kb) of the B6N NEERV (Figure 2A, bottom row). These data revealed that intergenic B6N NEERV loci possessed significantly increased activating and significantly reduced repressive histone modifications, suggesting that activation of B6N NEERV transcription occurs secondary to a primary failure of epigenetic silencing.

**Figure 2.**
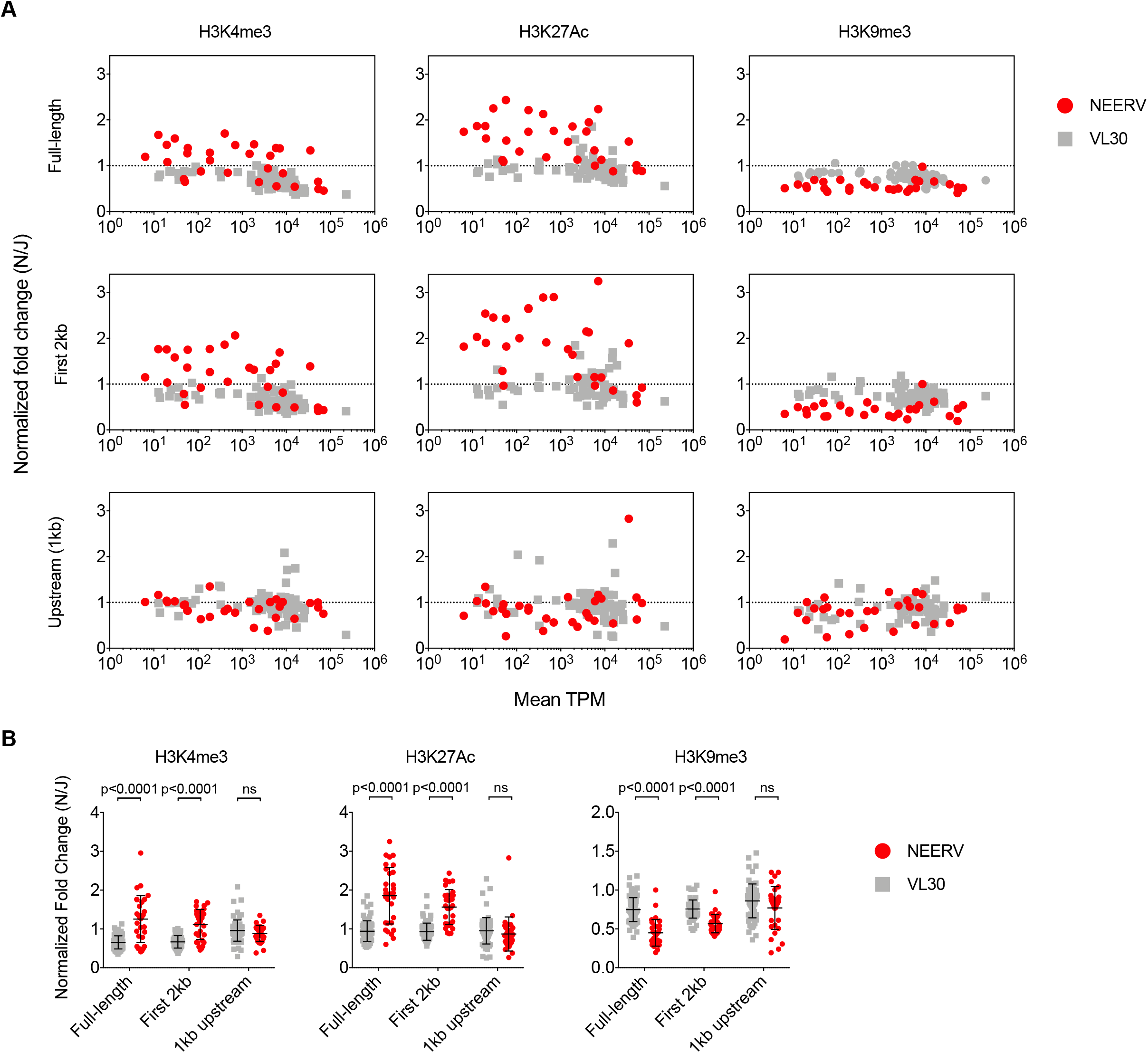
Intergenic NEERV loci are enriched for activating histone modifications and depleted of repressive histone modifications in BMDMs. **(A)** Plot of normalized fold change for each listed histone modification versus the mean expression level in B6N and B6J BMDMs in transcripts per million (TPM). Normalized fold change was calculated as: [(N_sum_ histone modification reads + 0.1)/(N_sum_ input reads + 0.1)]/ [(J_sum_ histone modification reads + 0.1)/(J_sum_ input reads + 0.1)]. This corresponds to: summation of the normalized ChIP-seq read counts across the full-length (top row), first 2kb (middle row), or 1kb immediately upstream (bottom row) of the NEERV (red) or VL30 (gray) loci for the histone modifications or input in B6N or B6J samples; addition of a pseudocount of 0.1 to all totals to avoid division by zero; division of the sums of the histone modifications by the sums of the input for the respective strain; and finally, division of the B6N-based value by the B6J-based value **(B)** Normalized fold changes plotted for each histone modification, with respect to each analyzed region as described above. Mean and standard deviation are plotted in black. Adjusted p-values were calculated for multiple t-tests comparing NEERV to VL30 for each histone mark across each region, corrected for the 9 independent hypotheses tested using the Holm-Sidák method with an alpha value of 0.05 for the entire family of comparisons.

### Recessive loss of proviral endogenous retrovirus silencing maps to a 1Mb deletion on chromosome 13

We next investigated the heredity of this phenotype by crossing B6N to B6J mice and evaluating the isogenic F1 generation. C57BL/6NJ F1 CD4 T lymphocytes expressed low NEERV envelope protein and mRNA levels (Figure 3A), demonstrating that the B6N phenotype of enhanced NEERV expression was recessive. Consistent with our ChIP-seq data, this suggested the existence of a NEERV repressor in B6J mice that is absent in B6N mice. To identify the genomic location that associates with the B6N phenotype, we performed a quantitative trait locus (QTL) analysis on 46 mice from the F2 C57BL/6NJ intercross. The mice were phenotyped for 5 parameters: surface ERV envelope expression on B, CD4 T and CD8 T cells; and total splenocyte *Xmv* and *Pmv* envelope mRNA expression (Figure S2A-C). Mice were genotyped at 150 SNPs that differentiate B6N and B6J substrains. We identified a single QTL locus on chromosome 13 significant for all 5 phenotypes (Figure 3B) and investigated nucleotide and structural variation across the predicted QTL interval from whole genome sequencing (WGS) data of B6N and B6J genomes (Figure S2D-F). In addition to identifying known (Simon et al., 2013) non-synonymous coding SNPs (Figure S2E), copy number variant analyses (Figure 3C & Figure S2F-G) supported by TaqMan Copy Number qPCR assays (Figure 3D) revealed a ~1Mb deletion within the QTL locus uniquely in the B6N genome (Figure 3E). No other structural variants were identified in the B6N genome that were not also present in the B6J genome. These findings indicated that loss of NEERV repression has occurred in the B6N substrain secondary to a deletion on chromosome 13.

**Figure 3.**
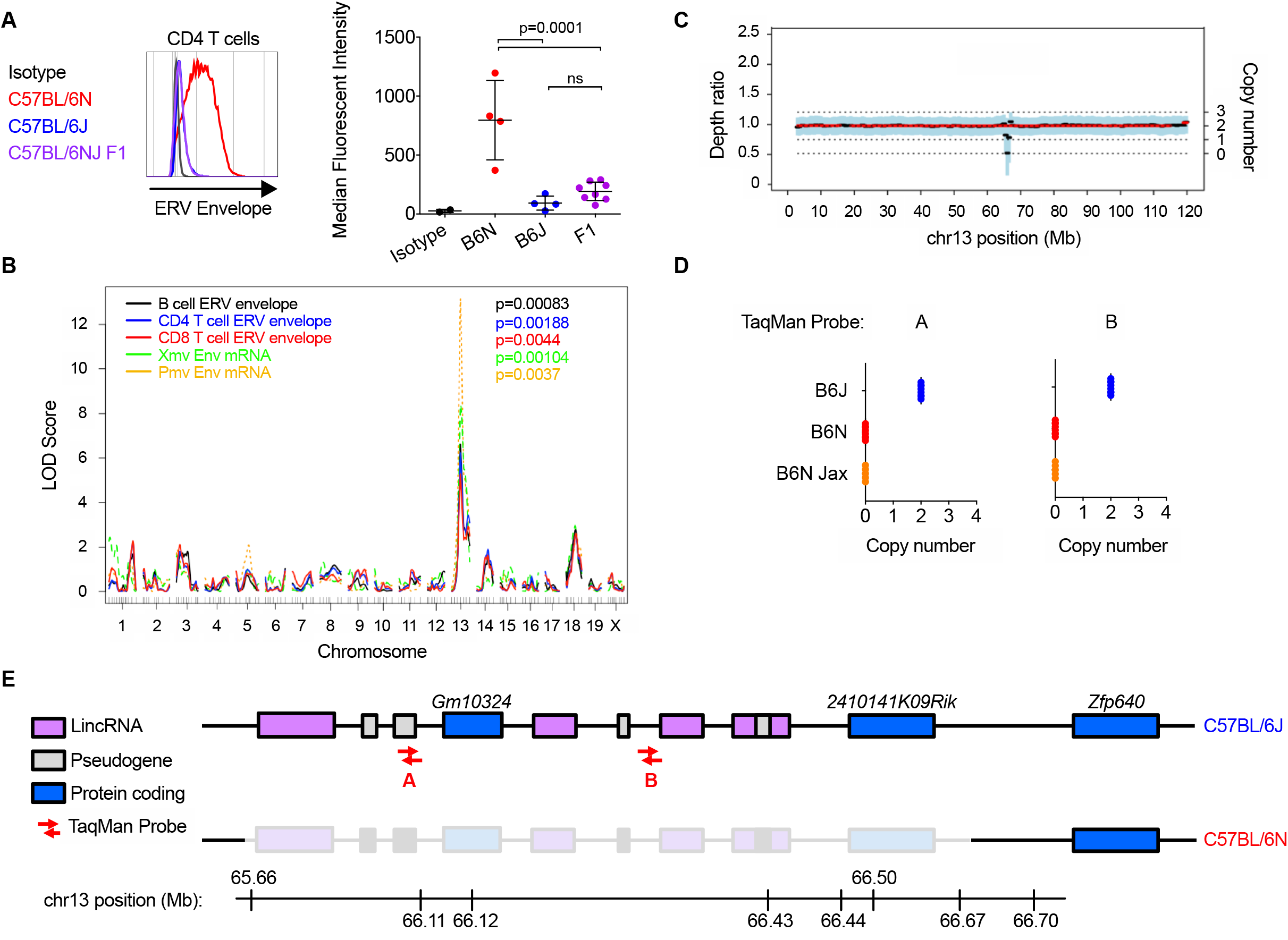
Recessive loss of proviral endogenous retrovirus silencing maps to a deletion in two KRAB-ZFP genes on chromosome 13. **(A)** Representative histogram or calculated MFI of ERV envelope protein expression detected via FACS on the surface of peripheral blood CD4^+^ T lymphocytes from adult mice. Each histogram or point represents an individual mouse. **(B)** Single-quantitative trait locus analysis from 46 F2 intercrossed C57BL/6NJ mice. The logarithm of the odds (LOD) score, comparing the hypothesis that there is a QTL at the marker to the null hypothesis that there is no QTL anywhere in the genome, is plotted for every SNP maker and imputed marker across the genome. **(C)** Sequenza estimates allele-specific copy number from paired tumor-normal sequencing data. Sequenza analysis comparing the B6J and B6N genomes identified a single region within the QTL interval in the B6N genome with a decrease in depth ratio and copy number. **(D)** TaqMan probes with unique binding sites within the region of interest were used to amplify product from B6J (Iwasaki colony) and B6N (Iwasaki and Jackson colonies) genomes. **(E)** The deleted region in the B6N genome spans several long intergenic non-coding RNAs & pseudogenes and 2 Krüppel-associated box zinc finger proteins. P-values in Figure 3A were calculated using one-way ANOVA with Šidák’s multiple comparisons test and an alpha value of 0.05. QTL P-values were calculated by performing 10,000 permutation tests to obtain a genome-wide distribution for the null hypothesis. See also Figure S2.

### Homozygous *2410141K09Rik^−/−^Gm10324^−/−^* mice fail to repress NEERV mRNA and protein expression

Within this deleted region of chromosome 13, there are 2 annotated coding genes, *2410141K09Rik* and *Gm10324* both of which are KRAB-ZFPs, 4 non-coding RNAs, and 3 pseudogenes (Figure 3E). To determine the gene(s) responsible for NEERV repression, we generated B6J mice deficient in one or multiple genes within the chromosome 13 region of interest using Clustered Regularly Interspaced Short Palindromic Repeats (CRISPR) technology (Figure 4A & Figure S3A). Due to the extremely repetitive nature of this chromosomal region, individual guide RNAs targeted multiple cut sites. Traditional PCR genotyping and sequencing was insufficient to confirm genetic deletions. Thus, we additionally used TaqMan probe amplification loss (Figure 4B) and whole genome 10x sequencing (Figure S3B-C). Of the 4 CRISPR-generated strains lacking portions of chromosome 13, only mice with a homozygous deletion of both *2410141K09Rik* and *Gm10324 (241Rik^−/−^Gm10324^−/−^)* were unable to repress NEERV by RT-qPCR (Figure 4C). By RNA-sequencing (Figure 4D), *241Rik-^/^-Gm10324^−/−^* phenocopied B6N mice, with concordance in both NEERV locus expression and magnitude of expression increase. This was also reflected in the levels of surface ERV envelope protein expression on lymphocytes (Figure 4E). Mice with a heterozygous deletion of these genes *(241Rik^-/+^ Gm10324^-/+^)* maintained B6J NEERV expression levels, confirming the haplosufficient nature of *241Rik* and *Gm10324* activity demonstrated by the C57BL/6NJ F1 mice (Figure 3A). Additionally, expression of non-NEERV RE was not increased (Figure S3D), validating the specificity for NEERV repression that was lost in the *241Rik^−/−^Gm10324^−/−^* mice. The expression of 8 cellular genes was also significantly increased more than two-fold in the *241Rik^−/−^Gm10324^−/−^* CD4 T cells (Figure 4D). Six of these genes directly overlap a NEERV LTR, suggesting that their increased expression may have resulted from a failure to silence the internal NEERV element. For example, *Camk2b* encodes for a neuronal protein kinase whose third intron contains an RLTR4_MM NEERV element. *Cam2kb* was one of the most significantly increased genes in both B6N and *241Rik^−/−^Gm10324^−/−^* CD4 T cells, suggesting that NEERV dysregulation, rather than substrain nucleotide differences (Simon et al., 2013), might have mediated this effect. Thus, the *241Rik^−/−^Gm10324^−/−^* phenotype indicated that one or both of these genes were responsible for silencing NEERV in the B6 genome and implicated NEERV dysregulation in increasing the expression of nearby cellular genes. We have therefore named *2410141K09Rik* and *Gm10324* suppressor of <u>N</u>E<u>ERV</u> 1 and 2 *(Snerv1* and *Snerv2),* respectively.

**Figure 4.**
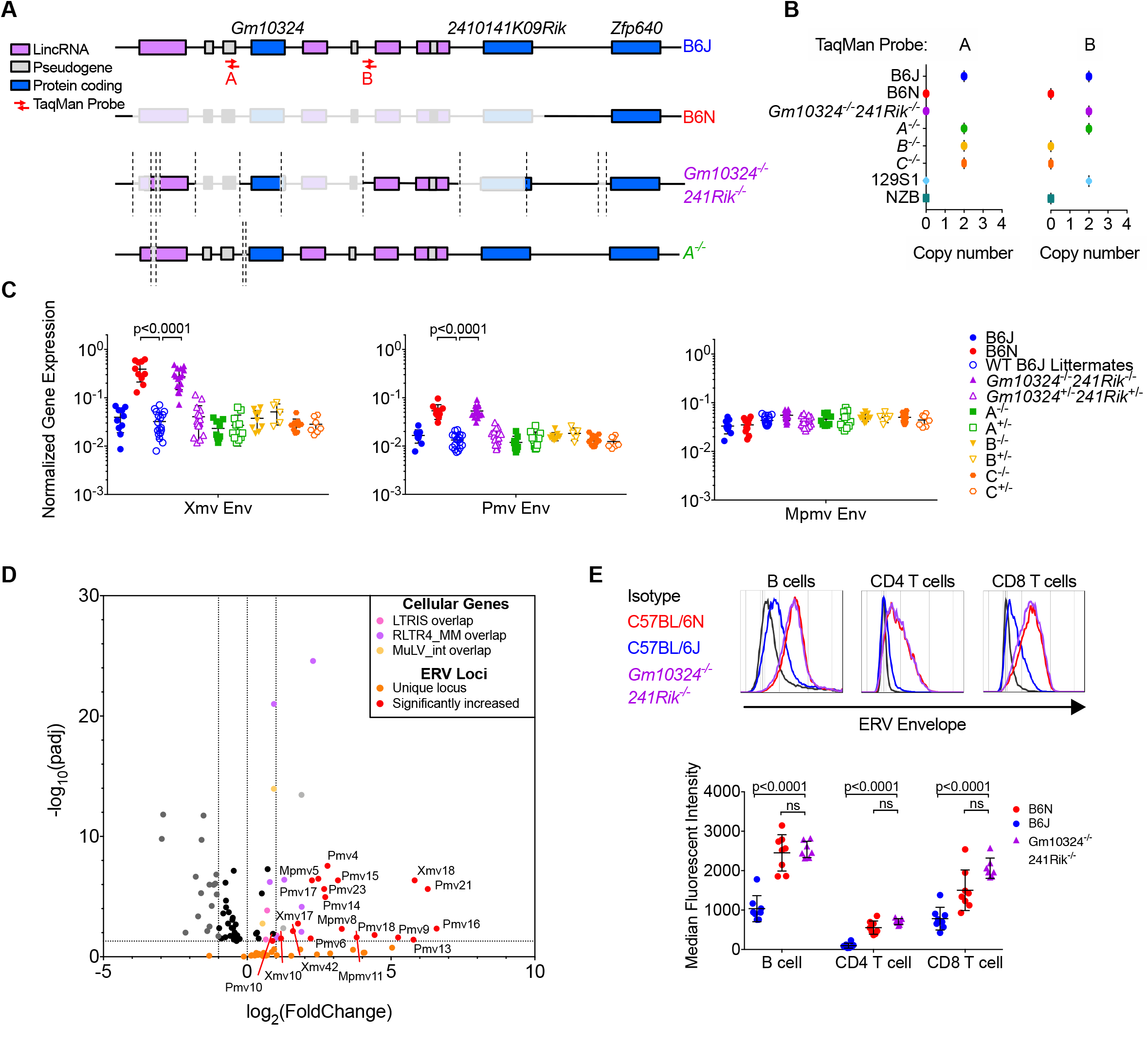
Homozygous *2410141K09Rik^−/−^Gm10324^−/−^* mice fail to repress NEERV mRNA and protein expression. **(A)** Schematic of chromosome 13 regions that were deleted in two of the B6J CRISPR-generated mice that were sequenced. **(B)** TaqMan probes with unique binding sites in the region of interest were used to amplify product from NZB, 129S1, B6N, B6J, and the CRISPR-generated mice (n=5 per group). **(C)** RT-qPCR of RNA from peripheral blood of WT and CRISPR-generated mice (n=5-18 per group) for Xmv, Pmv, and Mpmv envelope mRNA. Values were normalized to GAPDH expression. Listed are the significant adjusted p-values for multiple t-tests comparing all genotypes to the B6J WT littermate value for each gene, corrected for the 33 independent hypotheses tested in Figure 4 using the Holm-Sidák method with an alpha value of 0.05 for the entire family of comparisons. **(D)** Volcano plot of differentially expressed cellular genes & all 47 uniquely mappable ERV loci from mRNA sequencing of B6J and *241Rik^−/−^ Gm10324^−/−^* B6J CD4^+^ T cells. (E) Representative histogram and calculated MFI of ERV envelope protein expression detected via FACS on the surface of peripheral blood B cells, CD4^+^ T, and CD8^+^ T lymphocytes from adult B6J, B6N, and *241Rik^−/−^Gm10324^−/−^* mice. Each histogram or point represents an individual mouse. Adjusted p-values for Figure 4E were calculated for multiple t-tests comparing the *241Rik^−/−^Gm10324^−/−^* value to that of B6J (Figure 4E), corrected for the 33 independent hypotheses tested in Figure 4 using the Holm-Sidák method with an alpha value of 0.05 for the entire family of comparisons. Adjusted p-values in Figure 4D were calculated using DESeq2. See also Figure S3–S4.

The locus on chromosome 13 that contains *Snerv1* and *Snerv2* is remarkable for its extremely repetitive nature that necessitated WGS over PCR-based approaches for genotyping. Indeed, even stringent mapping criteria of commonly used sequencing alignment programs erroneously mapped a high proportion of low-confidence paired-end 150bp reads into this interval from genomes (such as B6N) in which this region was absent and therefore could not contribute reads (Figure S4A-C). Thus, alignments to and read counts for these genes using short-read sequencing technologies were not reliable. It is also of note that *Snerv1* and *Snerv2* were only expressed in early development (Figure S4C-E), preventing the detection of differential gene expression in somatic tissues.

### SNERV1, but not SNERV2, strongly recruits KAP1 and selectively binds to the glutamine-complementary primer binding site in the NEERV LTR

*Snerv1* and *Snerv2* genes both encode for KRAB-ZFPs, DNA-binding proteins that can bind to and silence RE through the recruitment of KAP1 and additional co-repressors (Ecco et al., 2017). Our *Snerv1/2^−/−^* phenotype suggested that one or both of these genes encode the KRAB-ZFP responsible for repression of NEERV. To determine if either of these proteins could interact with KAP1, because of the high homology to additional KRAB-ZFP loci and non-unique flanking sequences, both genes had to be codon-optimized and synthesized *de novo.* FLAG-tagged SNERV1 or SNERV2 was transiently overexpressed in HEK 293T cells to test the ability to immunoprecipitate with KAP1. FLAG-ZFP809 served as a positive control for KAP1 binding. Although both proteins were expressed to similar levels in nuclear extract, only FLAG-SNERV1, but not FLAG-SNERV2, strongly bound to KAP1 (Figure 5A).

**Figure 5.**
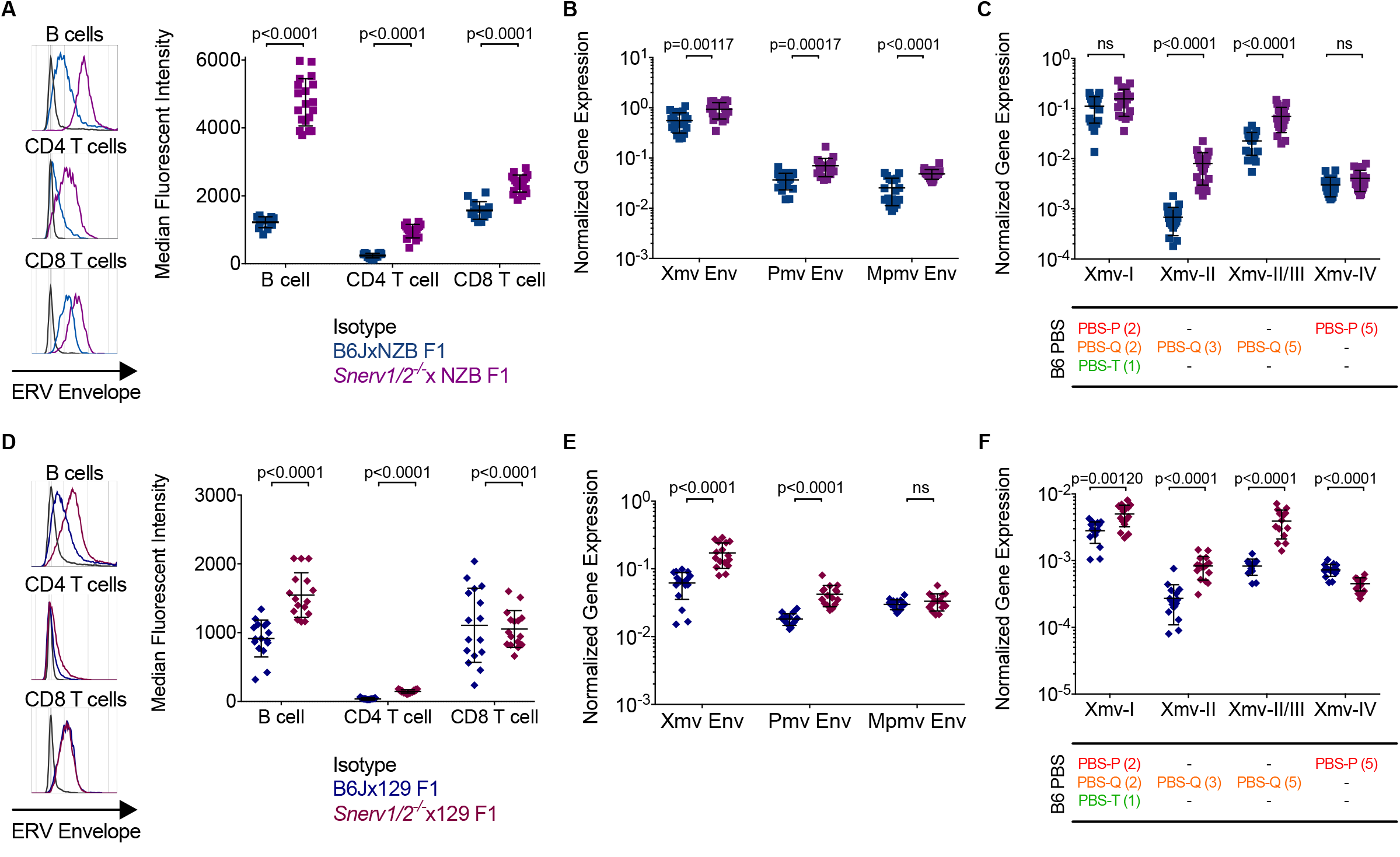
SNERV1, but not SNERV2, strongly recruits KAP1 and selectively binds to the glutamine-complementary primer binding site in the NEERV LTR. **(A)** Anti-FLAG and anti-KAP1 western blot of immunoprecipitated FLAG-ZFP from 293T nuclear lysate following transient overexpression of FLAG-ZFP809, FLAG-SNERV1, or FLAG-SNERV2. **(B)** Schematic of the ERV LTR and LTR-based oligos that were designed for use in DNA pulldown and electrophoretic mobility shift assays (EMSA). Primer binding sites of the LTR-based oligos are denoted by amino acid letter and color in (C)-(F). **(C)** DNA pulldown of 32bp biotinylated LTR oligos by recombinant GST-FLAG-SNERV1 or GST-FLAG-SNERV2. **(D)** DNA pulldown of 59bp biotinylated LTR oligos by recombinant GST-FLAG-SNERV1 or GST-FLAG-SNERV2. **(E)** EMSA of 54bp AlexaFluor488-labeled double-stranded LTR oligonucleotides (AF488-PBS) using no protein or 10ug of recombinant GST-FLAG-SNERV1 or GST-FLAG-SNERV2. **(F)** EMSA of 54bp AF488-PBS-Q using increasing amounts of recombinant GST-FLAG-SNERV1 or GST-FLAG-SNERV2. Competitor 59bp unlabeled PBS-Q and PBS-Q’ LTR oligonucleotides were used in lanes 5-6 and 10-11 in **(F)** in 10-fold excess. See also Figure S5.

The 5’ LTR of NEERV loci contain a GC-rich primer binding site (PBS) (Figure 5B) that is complementary to the 3’ end of cellular tRNAs and primes reverse transcription (Gilboa et al., 1979). ZFP809 binds to a proline-complementary PBS (PBS^Pro^) sequence, recruits KAP1, and represses ecotropic ERV transcription (Figure 5A) (Wolf and Goff, 2009). Forty-three of the 49 B6 NEERV sequences possess glutamine-complementary PBS (PBS^Gln^) (Table S1) (Jern et al., 2007). The expression of PBS^Gln^ NEERV was increased in B6N, suggesting that this substrain lacks the repressor that targets PBS^Gln^, and we hypothesized that SNERV1 and/or SNERV2 binds to the PBS^Gln^ sequence in the NEERV LTR. *Pmv15* is a PBS^Gln^-encoding NEERV whose expression is strongly repressed in B6J and highly increased in B6N CD4 T cells (Fig 1C). We generated 32bp DNA oligonucleotides spanning the *Pmv15* PBS sequence and the immediate downstream bases that improve binding of ZFP809 to its target PBS (Kempler et al., 1993). We also designed 54-59bp oligonucleotides that additionally include sequence from the upstream LTR. These Pmv15-based oligonucleotides were also modified to alternatively encode PBS^Pro^, PBS^Thr^, or PBS^Gln^ sequences (Figure 5B). We next produced purified recombinant GST-FLAG-tagged SNERV1 and SNERV2 proteins (Figure S5A) and performed DNA pull down and electrophoretic mobility shift assays (EMSA) to test the ability of SNERV1 and SNERV2 to bind these oligonucleotides. Recombinant SNERV1 strongly bound to the PBS^Gln^ oligonucleotide by DNA pull down (Figure 5C), and this binding was lost if PBS^Gln^ was replaced with PBS^Pro^, PBS^Thr^, or PBS^Phe^, or if the downstream motif was absent (Figure 5C-D). This suggested that both PBS^Gln^ and the downstream sequence were required for effective binding by SNERV1 to the NEERV LTR. In contrast to SNERV1, recombinant SNERV2 did not bind strongly to any of the PBS oligonucleotides. By EMSA, addition of recombinant SNERV1, but not SNERV2, caused significant slowing of the PBS^Gln^ oligonucleotide migration (Figure 5E-F). PBS^Pro^, PBS^Thr^, or PBS^Phe^ probes did not elicit this strong shift in signal (Figure 5E), which was competitively reduced upon the addition of excess unlabeled PBS^Gln^ oligonucleotide (Figure 5F). However, SNERV1 binding to the PBS^Gln^ probe was not competitively reduced by excess unlabeled PBS^Gln^’ oligonucleotide lacking the downstream 13bp motif (Figure 5F), providing further evidence that the PBS^Gln^ sequence and downstream motif were both required for specific binding of SNERV1 to the NEERV LTR. Accordingly, the presence of PBS^Gln^ was not sufficient for transcriptional repression, as the 10 intergenic VL30 loci that possess PBS^Gln^ were not differentially expressed upon loss of *Snerv1* and *Snerv2* (Figure S1C-D & Figure S5B). Additionally, these same loci were not enriched for H3K4me3 or H3K27Ac or depleted of H3K9me3 (Figure S5C). The 13bp sequence downstream of PBS^Gln^ VL30 differed at many residues from that found in the NEERV LTR (Table S1). Together, these data suggested that sequence within the NEERV LTR, in addition to PBS^Gln^, are required for specificity.

Although recombinant SNERV2 bound weakly to these oligonucleotides, SNERV2 was nevertheless similarly selective for the PBS^Gln^ sequence and requirement for the downstream motif (Figure 5E-F). *Snerv1* and *Snerv2* both encode a KRAB-A box and 14 and 19 zinc fingers, respectively. The 5 additional canonical zinc fingers of SNERV2 correspond to 140 amino acids, which produced 2 gaps in global pairwise alignment with SNERV1 (Figure S5D). However, the aligned amino acids of SNERV1 and SNERV2 shared 87% (404/464) identity and 93% (429/464) conservation. Given their genomic proximity and high degree of homology, *Snerv1* and *Snerv2* may have arisen through tandem duplication, thereby providing for inherent shared specificity for NEERV LTR PBS^Gln^. Collectively, while both proteins were selective for the PBS^Gln^ sequence in the NEERV LTR, only SNERV1 was capable of both stronger binding to the PBS^Gln^ sequence and better recruitment of KAP1.

### The NZB and 129 genomes fail to complement NEERV derepression in the *Snerv1/Z^−/−^* genome

Next, we examined the physiological relevance of SNERV loss in NEERV repression. NEERV expression is highly increased in NZB, NZW, and 129 strains and associates with lupus nephritis (Andrews, 1978; Baudino et al., 2008; Ito et al., 2013; Izui, 1979; Yoshiki et al., 1974). The *Sgp3* locus in NZB & NZW strains and the *Gv1* locus in 129 strains drive increased NEERV expression and are mapped by QTL analyses to similar large intervals on chromosome 13 that both include *Snerv1 and Snerv2* (Laporte et al., 2003; Oliver and Stoye, 1999). Loss of TaqMan probe amplification in proximity to both *Snerv* genes from NZB and 129 genomic DNA (Figure 4B) suggested that this interval might also be deleted in the NZB and 129 strains. Alignment quality of next-generation sequencing short reads across this chromosome 13 region from NZB and 129 genomes was extremely poor, exemplified by erratic read depth and copy number calls by both Sequenza and CNVnator (Figure S2F-G). We were unable to further clarify the structure of this region in the NZB genome using 10x WGS (Figure S6A-D) due to the highly tandemly repetitive nature of this genomic interval and the frequency of true SNPs and SVs that differentiate NZB from the B6J reference genome.

Therefore, to investigate if *Snerv1 and/or Snerv2* underlie the *Sgp3* and *Gv1* loci, we crossed *Snerv1/2^+l+^* (B6J wild-type) or *Snerv1/2^−/−^* females to NZB and 129 males to test for complementation. B6N/J F1 and *Snerv1/2^-/+^* mice both demonstrated that heterozygosity of these genes conferred haplosufficiency for NEERV repression. Therefore, if the *Sgp3* and *Gv1* loci do not involve these KRAB-ZFP genes, then both the NZB and 129 genomes will possess intact copies of these genes and will complement the *Snerv1/2^−/−^* genome to restore NEERV repression to levels exhibited by the B6J-NZB F1 cross.

While NZB and B6J mice possess many of the same Mpmv and Pmv loci, only 5 Xmv loci are shared (Frankel et al., 1992; Kihara et al., 2011). Additionally, NZB mice express high levels of Xmv mRNA from both constitutive and inducible Xmv loci (Elder et al., 1980). Unlike B6J mice, which predominantly express Xmv9, Xmv10, Xmv13, Xmv14 (all PBS^Gln^) and Xmv43 (PBS^Pro^) (Figure 1B & 1D), the highly expressed NZB Xmv NEERV encode PBS^Pro^ (Baudino et al., 2008; O’Neill et al., 1985). Xmv loci can be further subdivided into 4 subgroups, Xmv-I through Xmv-IV, whose transcripts can be amplified with subgroup-specific envelope primers. All B6 Xmv elements utilizing PBS^Pro^ belong to Xmv-I or Xmv-IV subgroups (Table S1), and the strongly expressed constitutive and inducible NZB Xmv loci are classified as Xmv-I (Baudino et al., 2008; Kihara et al., 2011). These PBS^Pro^-encoding Xmv-I and Xmv-IV elements should not be subject to SNERV-mediated repression, which is specific to PBS^Gln^.

Compared to B6JxNZB F1 pups, *Snerv1/2^−/−^xNZB* F1 pups expressed significantly higher levels of NEERV envelope protein on the surface of peripheral blood B cells, CD4 T cells, and CD8 T cells (Figure 6A). The expression of Xmv, Pmv, and Mpmv envelope mRNA was likewise significantly elevated in peripheral blood from these same mice, compared to B6JxNZB F1 controls (Figure 6B). Xmv-I expression was highly driven in both crosses, likely a consequence of the known constitutive PBS^Pro^ Xmv-I transcription that is characteristic of NZB mice. As expected, the expression of Xmv-I and Xmv-IV mRNA did not differ between the two crosses (Figure 6C). However, in contrast to their SNERV haplosufficient counterparts, *Snerv1/2^−/−^xNZB* F1 pups were unable to repress NEERV expression from PBS^Gln^-encoding Xmv-II and Xmv-III loci, leading to highly increased transcription from these loci (Figure 6C). These data indicated that SNERV proteins are required for B6J mice to repress Xmv, Pmv, and Mpmv loci in B6JxNZB F1 mice.

**Figure 6.**
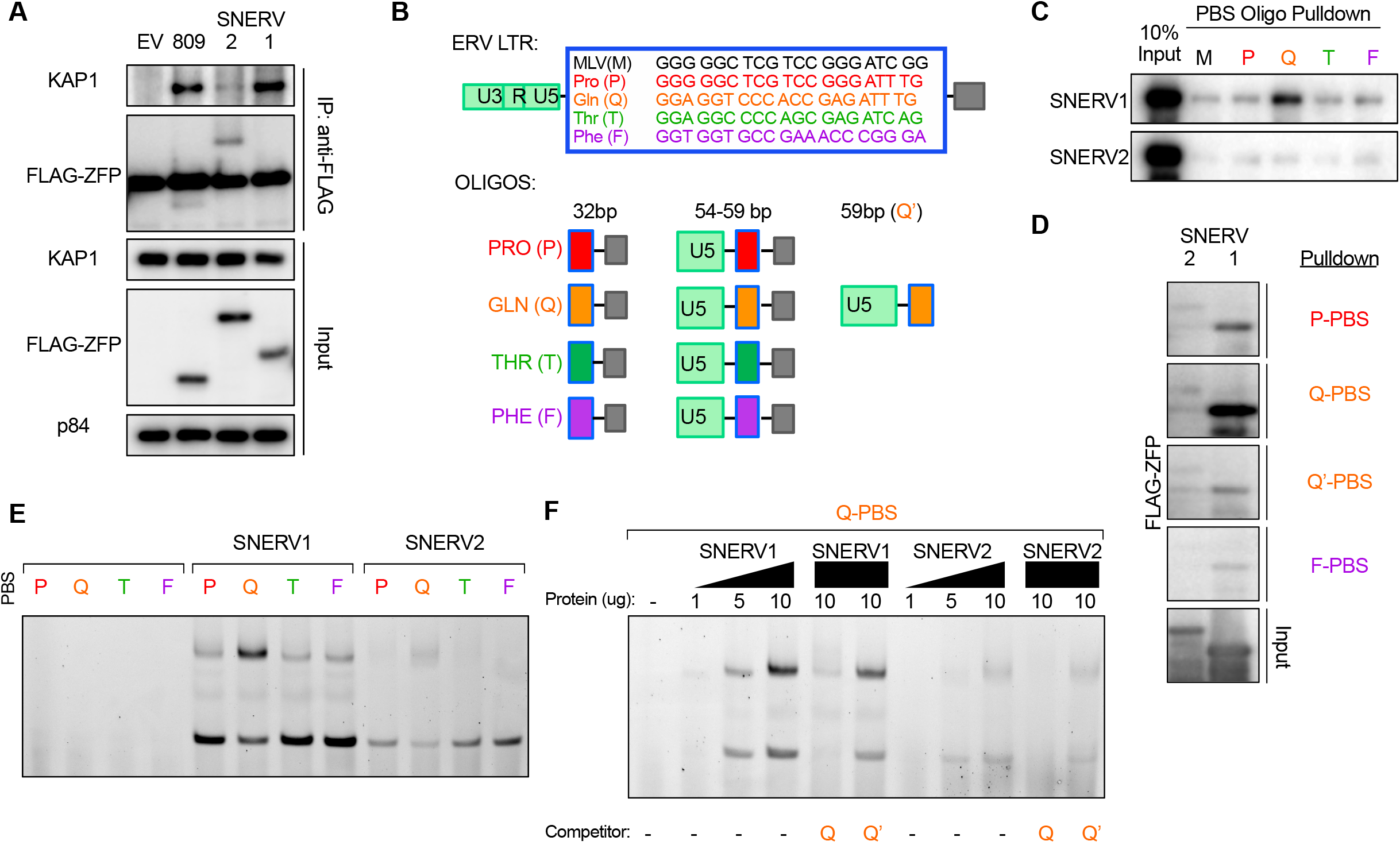
The NZB and 129 genomes do not complement the loss of NEERV silencing in the *Snerv1/2^−/−^* genome. **(A)**. Representative histogram and calculated MFI of ERV envelope protein expression detected via FACS on the surface of peripheral blood B cells, CD4^+^ T, and CD8^+^ T lymphocytes from adult B6JxNZB F1 and *Snerv1/2^−/−^xNZB* F1 mice. **(B)** RT-qPCR of RNA from peripheral blood from B6JxNZB F1 and *Snerv1/2^−/−^xNZB* F1 mice for Xmv, Pmv, and Mpmv envelope mRNA. **(C)** RT-qPCR of RNA from peripheral blood from B6JxNZB F1 and *Snerv1/2^−/−^xNZB* F1 mice for Xmv-I, Xmv-II, Xmv-II/III, and Xmv-IV mRNA expression. **(D)**. Representative histogram and calculated MFI of ERV envelope protein expression detected via FACS on the surface of peripheral blood B cells, CD4^+^ T, and CD8^+^ T lymphocytes from adult B6Jx129 F1 and Snerv1/2^−/−^x129 F1 mice. **(E)** RT-qPCR of RNA from peripheral blood from B6Jx129 F1 and Snerv1/2^−/−^x129 F1 mice for Xmv, Pmv, and Mpmv envelope mRNA. **(F)** RT-qPCR of RNA from peripheral blood from B6Jx129 F1 and Snerv1/2^−/−^x129 F1 mice for Xmv-I, Xmv-II, Xmv-II/III, and Xmv-IV mRNA expression. The PBS type(s) for mappable B6J Xmv loci are listed below their corresponding Xmv class, with the total number of loci in parentheses. Each histogram or point represents an individual mouse. Adjusted p-values were calculated for multiple t-tests comparing the *Snerv1/2*^−/−^-based F1 value to the B6J-based F1 value for each gene, corrected for the 20 independent hypotheses tested using the Holm-Sidák method with an alpha value of 0.05 for the entire family of comparisons. See also Figure S6.

Compared to B6J mice, 129 mice possess few Xmv loci and express near-undetectable levels of Xmv envelope transcripts (Baudino et al., 2008; O’Neill et al., 1986; Yoshinobu et al., 2009). As such, Xmv transcription in the B6Jx129 and *Snerv1/2^−/−^x129* F1 crosses arises largely from B6J loci. The *Gv1* locus controls Pmv, but not Mpmv, transcription in 129 mice (Oliver and Stoye, 1999). Compared to SNERV haplosufficient B6Jx129 F1 pups, the *Snerv1/2^−/−^x129* F1 pups expressed significantly higher levels of NEERV envelope protein on the surface of peripheral B cells and CD4 T cells (Figure 6D). Accordingly, Xmv and Pmv NEERV envelope mRNA was significantly increased in peripheral blood from these same mice, compared to B6Jx129 F1 controls (Figure 6E). As in *Snerv1/2^−/−^xNZB* mice, PBS^Gln^-encoding Xmv-II and Xmv-III envelope mRNA expression was significantly increased in *Snerv1/2^−/−^x129* mice (Figure 6F). Xmv-I transcription in *Snerv1/2^−/−^x129* F1 mice, which lack the high PBS^Pro^ Xmv-I expression of the NZB-based crosses that would otherwise mask its detection by RT-qPCR, was also significantly increased.

From the patterns of NEERV expression observed in the two sets of crosses, it is evident that control of NEERV expression is multifactorial: strain-specific locations and sequences of proviral NEERV; cell type-specific transcriptional programs that dictate which NEERV loci are in euchromatin; and strain- and cell-specific factors that regulate NEERV mRNA and protein synthesis and degradation. Yet unlike B6JxNZB and B6Jx129 F1 mice, both *Snerv1/2^−/−^xNZB* and *Snerv1/2^−/−^x1*29 F1 mice were unable to repress NEERV. These data demonstrated that functional SNERV are absent in both NZB and 129 genomes. Although we could not rule out an effect from nearby intergenic deletions that are also present in the *Snerv1/2^−/−^* genome (Figure 4A), two similar non-coding deletions, including that in the *Platr2* pseudogene, were present in the *A^−/−^* genome and did not give rise to increased NEERV expression. This suggested that such non-coding deletions do not impact the function of *Snerv1* or *Snerv2* or otherwise modulate expression of NEERV. Thus, the failure of the NZB and 129 genomes to complement the loss of these genes implicates defective SNERV as both the *Sgp3* and *Gv1* loci.

### Human ERV LTR elements are elevated in the blood of patients with SLE and identification of putative HERV suppressing KRAB-ZFPs

Our data support a role for KRAB-ZFP-mediated loss of NEERV suppression in murine lupus pathogenesis. To investigate the relevance of these findings to human SLE, we interrogated publicly available RNAseq data of whole blood from SLE patients and healthy controls (Hung et al., 2015; Kalunian et al., 2015) for HERV expression, using the RepEnrich algorithm to quantitate read counts for HERV LTR families and subfamilies. A number of LTR subfamilies were significantly elevated in SLE blood compared with healthy controls (Figure 7A). While some SLE patients expressed low levels of ERVs, comparable to healthy controls, the majority of SLE patients expressed elevated levels of all LTR subfamilies, compared with healthy controls (Figure 7B). SLE patients expressed elevated levels of ERVL-MaLR, ERV1, and ERVL, which represent class I gammaretroviruses (ERV1) and class III spuma-like retroviruses (ERVL-MaLR and ERVL) (Figure 7C).

**Figure 7.**
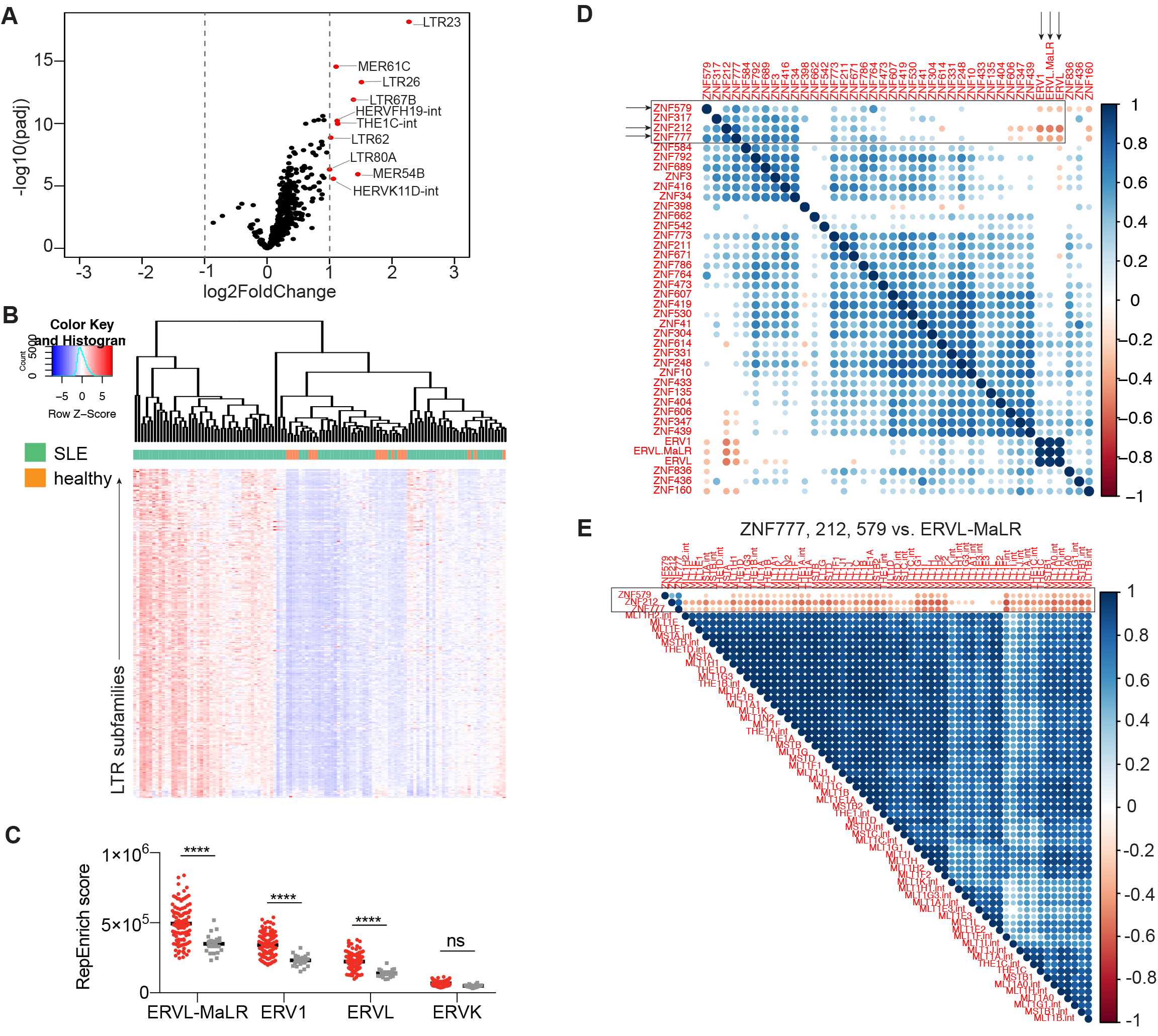
HERV LTR elements are elevated in the blood of patients with SLE and identification of putative HERV-suppressing KRAB-ZFPs. RNA sequencing data from whole blood of SLE patients (n=99) and healthy controls (n=18) were used to perform RepEnrich and DESeq2 analyses to quantify expression of LTR elements and cellular genes, respectively. **(A)** Volcano plot of significantly elevated LTR subfamilies in the blood of SLE patients versus healthy controls. LTR subfamilies indicated in red are log_2_(Fold Change) > 1 and padj < 0.05 in SLE patients versus healthy controls. **(B)** Heatmap of all LTR subfamilies that are significantly differentially expressed in SLE patients compared with healthy controls (padj < 0.05, n=316). Hierarchical clustering of patients was performed based on Euclidean distance. **(C)** The sum of all reads that belong to each indicated LTR families was graphed per individual. Two-way ANOVA was performed to calculate statistical significance. ****, p < 0.0001; ns, not significant. **(D-E)** Spearman correlation was calculated between all of the repressed KRAB-ZFPs and the sum of RepEnrich scores for the significantly elevated LTR families **(D)**, and LTR subfamilies that belong to the ERVL-MaLR and ZNF777, ZNF212, and ZNF579 **(E)** among SLE patients. The correlation plot represents Spearman r values and displays only correlations that were p < 0.05. Blank indicates not significant. See also Figure S7.

In an effort to identify potential KRAB-ZFPs that could function as suppressors of HERV, that may be dysfunctional in SLE patients, we performed a Spearman correlation analysis between the 38 KRAB-ZFP genes that were significantly repressed in SLE patients (Figure S7A) and the sum of RepEnrich scores for ERVL-MaLR, ERV1, and ERVL families. Three KRAB-ZFPs were significantly negatively correlated with all 3 LTR families: ZNF777, ZNF579, and ZNF212 (Figure 7D). When expression of these KRAB-ZFPs was correlated with the expression of all of the LTR subfamilies within each of the LTR families, these KRAB-ZFPs and most of the HERV subfamilies were consistently and significantly negatively correlated (Figure 7E and Figure S7B-C). Thus, analogous to SNERV, these KRAB-ZFPs may function as suppressors of HERV, and decreased expression of these KRAB-ZFPs in SLE patients may contribute to the elevated HERV expression that was observed.

## Discussion

Our study identified *Snerv1* and *Snerv2,* encoding KRAB-ZFPs responsible for NEERV repression in multiple inbred mouse strains. SNERV targeted the PBS^Gln^ sequence within the NEERV LTR and recruited KAP1 protein to promote formation of heterochromatin at NEERV loci. Germline homozygous deletion of two KRAB-ZFP, *Snerv1* and *Snerv2,* increased activating chromatin modifications, transcription, and expression of protein from NEERV loci. F1 crosses of lupus-prone NZB and 129 mice to *Snerv1/2^−/−^* mice were unable to rescue defective NEERV repression, thus mapping the lupus-associated *Sgp3* and *Gv1* loci to *Snerv1* and *Snerv2* and demonstrating that loss of SNERV drove overexpression of the lupus autoantigen, gp70. Similar to how SNERV loss and resultant NEERV dysregulation are a hallmark of spontaneous lupus disease in mice, global increases in HERV family and subfamily expression was a salient transcriptional feature of SLE disease in humans. Antibodies against specific HERV antigens are present in SLE patients (Bengtsson et al., 1996; Blomberg et al., 1994; Nelson et al., 2014; Perl et al., 1995), yet it not known how HERV antigen overproduction results in this loss of tolerance. Having identified SNERV as the KRAB-ZFPs targeting NEERV in lupus-prone NZB and 129 mice, it will now be possible to define the contribution of ERV to lupus nephritis pathogenesis and test how ERV misregulation mediates loss of tolerance in murine and human lupus disease.

Restoration of SNERV1 and SNERV2 to the germline represses NEERV loci and prevents gp70 overproduction in lupus-prone mice. Generation of *Snerv1/2-competent* NZB and NZW will permit targeted approaches to manipulate the gp70 phenotype *in vivo*, and can be used to conclusively test the requirement for dysregulated NEERV in the pathogenesis of lupus. While *Snerv1 and Snerv2* are in epistasis with additional susceptibility loci that enhance disease in models of spontaneous lupus (Celhar and Fairhurst, 2017; Crampton et al., 2014; Morel, 2010), such experiments will elucidate the connection between ERV misregulation and lupus pathogenesis. Using Snerv1/2-competent lupus-prone mice, it will be possible to rigorously test how tolerance is lost in the setting of high NEERV autoantigen production, how anti-NEERV autoantibodies are induced, and how NEERV dysregulation itself contributes to lupus severity. Establishing the precise role of NEERV autoantigen overexpression in murine nephritis will likewise contribute to our general understanding of how loss of tolerance and autoantibody production occur in autoimmunity.

NZB-and NZW-based lupus models are widely used in pre-clinical drug efficacy trials, as they recapitulate more clinical features of human SLE than other mouse strains (Celhar and Fairhurst, 2017; Li et al., 2017). Of the few drugs approved for treatment of SLE by the FDA, essentially all—systemic immunosuppressants, antimalarials, anti-BAFF, anti-CD20, anti-CTLA-4, interferon-alpha blockade, and toll-like receptor agonists—were tested pre-clinically in NZB/W models (Celhar and Fairhurst, 2017). Clarifying the role of NEERV in disease progression will shape how pre-clinical testing for lupus nephritis proceeds and whether it may be feasible to pursue the development of therapeutics that target ERV. In these ways, our identification of *Snerv1* and *Snerv2* and the mechanism of NEERV repression has many applications to the study of both human lupus pathogenesis and treatment.

KRAB-ZFP that target RE tend to emerge following genomic invasion by the retrovirus that they target. While KRAB-ZFP are broadly conserved in mammals, a large subset of rodent KRAB-ZFP are specific to the order Rodentia (Imbeault et al., 2017). With full-length retrovirus architecture and intact open reading frames, NEERV are among the more recently endogenized murine RE (Tomonaga and Coffin, 1998). While *Snerv1* and *Snerv2* have orthologs in the rat and hamster genomes, none are found in the human genome. We therefore posit that these genes emerged in the last common ancestor of mice, rats, and hamster shortly following the invasion of its genome by the MLV-type retrovirus that it targets. Yet just as the presence of a PBS is conserved across retroelements, PBS targeting is highly conserved across different KRAB-ZFP, regardless of their target species (Ecco et al., 2016; Wolf and Goff, 2007; Wolf et al., 2008). This suggests that the mechanism of PBS^Gln^ targeting may very well be conserved in a different human KRAB-ZFP/HERV pairing. Three human KRAB-ZFP were identified whose expression was significantly repressed in SLE patients and whose levels were significantly anticorrelated with increased HERV expression in SLE patients. Although our current study does not provide functional evidence, investigating polymorphisms in and near these 3 KRAB-ZFP genes, and epigenetic regulation of these KRAB-ZFP, in SLE and healthy cohorts could prove informative.

Thus, with broad implications for human SLE and autoimmunity, identification of *Snerv1* and *Snerv2* and their mechanism of NEERV repression will permit interrogation of the association between NEERV overexpression and murine lupus pathogenesis. Our finding that *Snerv1 and Snerv2* underlie the lupus-associated *Sgp3* and *Gv1* loci will provide for the development of new genetic tools in the study of murine lupus, and the *in vivo* demonstration that *Snerv1/2^−/−^* yields misregulation of the NEERV gp70 autoantigen provides a framework for improving our understanding of HERV misregulation in human SLE.

## STAR Methods

### Contact for Reagent and Resource Sharing

Further information and requests for resources and reagents should be directed to and will be fulfilled by the Corresponding Author, Akiko Iwasaki.

### Experimental Model and Subject Details

#### Mice

C57BL/6N mice were obtained from Charles Rivers and bred in-house. C57BL/6NJ (stock #005304), C57BL/6J (stock #000664), 129SvImJ (stock #002448), and NZB/BlNJ (stock #000684) mice were obtained from Jackson Laboratories. C57BL/6J mice were bred-in house. Mice were housed in SPF conditions and care was provided in accordance with Yale University IACUC guidelines (protocol #10365).

#### Primary Cultures Peripheral blood & splenocyte isolation

Mice were anesthetized and blood was obtained via retro-orbital bleed. Mice were sacrificed using CO_2_ inhalation followed by cervical dislocation in accordance with IACUC protocols and NIH guidelines. Blood was collected with heparinized Natelson tubes (Fisher Scientific) into 8mM EDTA in PBS. Red blood cells were lysed with ACK lysis buffer (150mM NH_4_Cl, 1 M KHCO_3_, 0.1mM EDTA, pH 7.4) and cells were washed twice with PBS before addition of RLT buffer (Qiagen). Spleens were dissociated through a 40μm filter in RPMI media (Gibco), red blood cells were lysed as above, and splenocytes were washed and passed through a 70μm filter prior to counting. Naïve or bulk CD4 T lymphocytes were using negative selection with the EasySep Mouse Naïve CD4 T cell isolation kit or the EasySep Mouse CD4 T cell isolation kit (StemCell). For RNA or DNA+RNA isolation, samples in RLT were spun through QIAShredder columns (Qiagen). All samples were stored at -80 prior to RNA isolation.

#### Total bone marrow isolation and bone marrow-derived macrophage generation

Bone marrow-derived macrophages were isolated as described by harvesting femurs and tibias from mice, removing all muscle tissue, and crushing the bones with a mortar and pestle in RPMI to release the marrow. The bone marrow suspension was homogenized by pipetting and then passed through a 70um filter into a 15mL conical. Red blood cells were lysed with ACK lysis buffer and cells were washed twice with RPMI before resuspending in complete RPMI and counting cells. Bone marrow cells were cultured in complete RPMI media supplemented with 50ng/mL recombinant macrophage colony stimulating factor (BioLegend) or with 30% L929-conditioned media. Cells were cultured for 7 days before lysis in RLT buffer for RNA isolation or subsequent chromatin immunoprecipitation. Media was replaced every two days for the duration of culture.

#### Murine embryonic fibroblast generation

Pregnant C57BL/6N and C57BL/6J female mice were sacrificed and E14.5 embryos were harvested and processed by first removing heads and livers. The remaining tissue was placed in a petri dish with 2.5mL of 0.05% Trypsin-0.5mM EDTA-PBS and minced with a scissors. The minced tissue was transferred to a 15mL conical, incubated in a 37-degree Celsius shaking water bath for 40min, and dissociated by adding 3mL of DMEM with 10% FBS and pipetting vigorously. The isolated cells were filtered through a 70um filter, resuspended in 15mL of DMEM with 10% FBS, and plated in a 20cm tissue culture dish. The cells were grown to confluency prior to freezing down (considered passage 1). MEFs were grown in 10cm tissue culture plates in 10%FBS in DMEM with 1x penicillin-streptomycin and isolated for RNA at passage 2. Cells were expanded to passage 4 and inactivated with Cesium-137 irradiation for use as ESC feeders.

#### Embryonic stem cell culture

C57BL/6N and C57BL/6J embryonic stem cells were obtained from Riken Institute (cell lines AES0143 and AES0144). Cells were cultured by pre-coating the tissue culture vessel with 1% gelatin (Stem Cell Technologies 7903), and then plating the embryonic stem cells along with irradiated C57BL/6J MEFs in maintenance media composed of Knockout DMEM (Gibco), 1x GlutaMax (Gibco), 1x non-essential amino acids (Gibco), 100uM beta-mercaptoethanol, 1x penicillin-streptomycin, and 1000U/mL rmLIF (Millipore ESG1107), supplemented with 20% Knock-Out Serum Replacement (Gibco). Confluent cells were split every 2-3 days for maintenance of the cell line, using 0.25% Trypsin-EDTA (Gibco) to dissociate the cells from the plate. Prior to cell harvest for RNA isolation, cells were plated off of feeders, on tissue culture plates pre-coated with gelatin. Cells were harvested 48 hours later into RLT buffer for RNA isolation.

#### Oocyte Isolation

All injections and oocyte isolations were performed by the Yale Genome Editing Center. Mature denuded oocytes were isolated as described (Guzeloglu-Kayisli et al., 2012) from 6 C57BL/6N and 6 C57BL/6J 3-week-old female mice. Mice were injected intraperitoneally (IP) with 5 IU of pregnant mare serum gonadotropin (PMSG) to induce superovulation. Forty-eight hours later, mice were injected IP with 5 IU of human chorionic gonadotropin (hCG), and mice were sacrificed by cervical dislocation 14hr after this second injection. Ovaries were harvested and the placed into a 60mm petri dish containing pre-warmed M2 medium (Millipore MR-015-D). After dissociating oocytes from the follicles, cumulus cells were detached from the oocytes by addition of 0.3mg/mL hyaluronidase and repeated pipetting of the cumulus-oocyte complexes using a capillary tube microinjection pipette. Oocytes were then washed three times by transferring the cells into new droplets of media and oocytes were counted under the microscope and then transferred into RLT buffer (Qiagen) supplemented with beta-mercaptoethanol. Samples were vortexed and then frozen at -80 degrees Celsius. Immature denuded MI prophase-arrested (germinal vesicle) oocytes were isolated as described (Guzeloglu-Kayisli et al., 2012) from 4 C57BL/6N and 4 C57BL/6J 3-week-old female mice 44hr after IP injection with PMSG and using culture media containing 10uM milrinone to ensure metaphase arrest (Stein and Schindler, 2011).

#### Flow cytometry

0.5-1 million splenocytes were plated in a 96-well round-bottom dish and stained with LIVE/DEAD™ Fixable Aqua Dead Cell Stain (ThermoFisher) followed by F_c_ block (clone 2.4G2). To stain for ecotropic and non-ecotropic ERV envelope protein, cells were incubated with hybridoma 83A25 supernatant (Leonard Evans, Rocky Mountain Laboratories) or rat IgG_2A_ isotype control, followed by mouse anti-rat IgG_2_A-biotin and streptavidin-PE-Dazzle594 (BioLegend). Cell surface markers were stained with anti-mouse CD3-APC, B220-BV605, CD4-APC-Cy7, and CD8-FITC (BioLegend). All incubations were performed at a final volume of 30μL for 15-20min at 4 degrees Celsius. Flow cytometry data was analyzed with FlowJo.

#### Reverse transcription-quantitative polymerase chain reaction (RT-qPCR)

RNA was isolated from total splenocytes or peripheral blood using either the RNeasy Kit or the AllPrep DNA/RNA Kit (Qiagen). cDNA was synthesized using iScript™ cDNA Synthesis Kit (BioRad). Quantitative PCR was performed using iTaq™ Universal SYBR^®^ Green Supermix (BioRad) in 10ul reactions in triplicate using 5-30ng of cDNA per reaction. Primers were used at a final concentration of 0.225μM and sequences are listed in Table S2.

#### CD4^+^ T-cell RNA library preparation & sequencing

RNA was isolated from naïve and bulk CD4 T cells using the RNeasy Kit and 500ng was used for paired-end library generation with the Illumina TruSeq RNA Library Prep Kit (naïve) or the NEB Ultra RNA Library Prep Kit (bulk). Libraries were run on a NextSeq500 to generate 2×75bp or 2×150bp reads.

#### Oocyte RNA library preparation & sequencing

Oocyte isolations were performed twice to generate biological duplicates, and oocytes were pooled into RLT buffer and stored at -80 degrees Celsius prior to RNA isolation with the RNAeasy Micro Kit (Qiagen). The number of pooled oocytes ranged from 260-310 (immature) and 133-195 (mature). Due to the low number of cells, sequencing libraries were generated using a modified single cell 96-well plate-based protocol (Haber et al., 2017). Purified RNA was captured using 2.2X RNAClean XP beads (Agencourt). RNA+beads were incubated on a magnet plate (Alpaqua Magnum FLX) and washed twice with 80% ethanol and air dried. Dried beads were resuspended in 8ul of Master Mix 1 (2.5 μM 3’ RT primer, 2.5 mM dNTPs (ThermoFisher), 1 unit RNAase inhibitor (Takara)) and incubated at 72° C for 3 minutes, after which the plate was immediately placed on ice for 1 minute to denature the RNA. After this incubation, 14ul of Master Mix 2 (1.4X Maxima RNase H-minus RT buffer (Thermo-Fisher), 1.4 M Betaine (Sigma), 12.9 mM MgCl2 (Sigma), 1.4 μM Template Switching Oligo, 1.4 units RNAase inhibitor (Takara), 2.9 units Maxima RNase H-minus RT (Thermo-Fisher)) was added to each well. The plate was then incubated at 50° C for 90 minutes followed by 85° C for 5 minutes for reverse transcription. Following reverse transcription, 28 ul of Master Mix 3 (0.4 μM ISPCR primer, 1.8X Kapa HiFi HotStart ReadyMix) was added to each well. cDNA was them amplified for 12 cycles (98° C for 3 minutes followed by 12 cycles of 98° C for 15 seconds, 67° C for 20 seconds, 72° C for 6 minutes followed by 72° C for 5 minutes). Amplified cDNA was purified using 0.7X AMPure XP beads (Agencourt) and washed twice with 70% ethanol and air dried. Dried beads were resuspended in 40 ul of TE and 35ul of DNA was transferred into a new well. DNA was quantified using a Qubit dsDNA HS kit and DNA was normalized to 0.2 ng/ul. 5ul of normalized DNA was used to generate RNA-seq libraries using the Nextera XT kit (Illumina). Primers used: 3’ RT primer 5-AAGCAGTGGTATCAACGCAGAGTACT30VN-3’ (Sigma); Template Switching Oligo 5’-AAGCAGTGGTATCAACGCAGAGTACATrGrG+G-3’ (Exiqon); ISPCR 5’-AAGCAGTGGTATCAACGCAGAGT-3’ (Sigma).

#### RNA sequencing analysis

RNA sequencing data from naïve CD4 T cells, bulk CD4 T cells, BMDMs, mature and immature oocytes, and public data from SRP018525 (Xue et al., 2013) and SRP059745 (Veselovska et al., 2015) were analyzed as described below.

##### Cellular Genes

The raw reads of RNA-seq experiments were trimmed of sequencing adaptors and low-quality regions by Btrim (Kong, 2011). The trimmed reads were mapped to mouse genome (GRCm38; mm10) by Tophat2 (Kim et al., 2013). After the counts are collected, the differential expression analysis was performed using DEseq2 (Love et al., 2014), which calculated the fold changes and adjusted p-values.

##### ERV (mapped to genome)

The Illumina reads were first trimmed by Btrim (Kong, 2011) to remove sequencing adaptors and low-quality regions. The trimmed reads were mapped to the mouse genome (GRCm38) using BWA-mem (Li and Durbin, 2010) with default parameters. The unmapped reads were filtered out using SAMtools (Li et al., 2009) and the mapped reads in SAM format were further processed as the following. The CIGAR field in the SAM file was used to check the number of hard or soft clipping. If the ratio of sum of hard and soft clipping to the length of the read was greater than or equal to 0.02, then the read was discarded. The remaining reads were checked for the field of edit distance compared to the locus reference (NM field). If the ratio of the edit distance to the sequence read length was greater or equal to 0.02, the read was discarded. Finally, the difference between the alignment score (field AS) and the suboptimal alignment score (field XS) was compared. If the difference was less than 5, the read was discarded. The SAM file that contained the mapped reads that pass the filtering steps described above was converted to a BAM file using SAMtools. This BAM file, together with the file that contains the ERV coordinates in the mouse genome (GRCm38) in bed format, was used as input to count the read mapping in each ERV locus by BEDTools (Quinlan and Hall, 2010). The read counts were normalized by the size factors obtained from the cellular genes of the same sample, calculated using the DESeq2 normalization method. ERV were also mapped to ERV sequences using a reference sequence containing the ERV sequences during the mapping stage, instead of the reference mouse genome.

##### Analysis of repetitive element enrichment (RepEnrich)

The raw reads of RNA-seq experiments were trimmed of sequencing adaptors and low quality regions by Btrim (Kong, 2011). The trimmed reads were first mapped to the mouse genome (mm10) using Bowtie (Langmead et al., 2009) with options that only allow unique alignments. The reads that mapped to multiple locations were written to separate files. The SAM output file from Bowtie that contained uniquely mapped reads was converted to a BAM file with Samtools and sorted. The sorted BAM file, the file that contained the reads that mapped to multiple locations, and the BED file that contained annotation of target repetitive elements (downloaded from the RepeatMasker track from UCSC genome table browser) were used as input for RepEnrich (Criscione et al., 2014). RepEnrich first tested the uniquely mapped reads for overlap with annotated repetitive elements. Then, RNA-seq reads mapping to multiple locations were mapped to repetitive element pseudo-genomes that represent all annotated genomic instances of repeat sub-families. If a read mapped to a single repeat sub-family pseudo-genome, it was counted once within that repeat sub-family, while reads mapping to multiple repeat sub-family pseudo-genomes were assigned a value equal to the inverse of the number of repeat subfamilies aligned. The repeat element sub-family enrichment was equal to the sum of these two numbers rounded to the nearest integer.

#### Chromatin immunoprecipitation & sequencing

B6N and B6J BMDMs were crosslinked with 1% formaldehyde (EMS) in 15 cm TC plates for 10 minutes with gentle shaking at room temperature. The reaction was quenched by adding 125 mM final concentration of glycine for another 5 minutes with shaking at room temperature. The cells were washed 3 times with cold PBS and scraped with 10 mls of PBS into 50 ml conical tubes. The cells were centrifuged for 5 minutes at 1,500 rpm at 4 degrees and resuspended in 1 ml cell lysis buffer per 15 cm plate (10 mM Hepes pH 7.3, 85 mM KCl, 1 mM EDTA, 0.5% IGEPAL CA-630, 1x protease inhibitors (ThermoFisher Halt) and incubated on ice for 5 minutes. The lysate was centrifuged for 5 minutes at 4,000 rpm at 4 degrees, the supernatant was removed, and the pellet resuspended in 0.3 ml of nuclear lysis buffer per 15 cm plate (10mM Tris pH 8.0, 0.5% N-lauroylsarcosine, 0.1% sodium deoxycholate, 100 mM NaCl, 1 mM EDTA, 0.5 mM EGTA, 1x protease inhibitors (ThermoFisher Halt)) and transferred to a 1.5 ml Bioruptor Plus TPX microtube (Diagenode). The nuclear lysate was sonicated for 3 rounds of 15 cycles of 30 seconds on/30 seconds off on high power (Diagenode Biorupter Plus). After sonication the chromatin was centrifuged for 15 minutes at max speed at 4 degrees and the supernatant transferred to a new tube. Triton X-100 was added to 1% final concentration. 0.7 mls of ChIP dilution buffer was added per 15 cm plate (20 mM Tris pH 7.5, 0.5% Triton X-100, 100 mM NaCl, 1 mM EDTA, 1x protease inhibitor (ThermoFisher Halt)) to the sonicated chromatin. Input was removed from diluted chromatin and frozen at -20 degrees. The remaining diluted chromatin was split into low binding tubes (one tube per antibody) and 5 ug of antibody added overnight and rotated at 4 degrees. Approximately 10 million BMDMs were used per ChIP, antibodies used were anti-H3K4me3 (Abcam ab8580), anti-H3K9me3 (Abcam ab8898), and anti-H3K27Ac (Active Motif 39133). The next day protein G Dynabeads (ThermoFisher 10004D) were washed 3X with PBS + 0.5% BSA (50ul of Dynabeads were used per IP). A Dynal magnet (Invitrogen) was used for Dynabeads washing and eluting steps. After washing, 50ul of dynabeads in PBS+BSA were added to each overnight tube containing the chromatin and antibody and rotated at 4 degrees for 3 hours. After rotation the beads were washed 3 times with low-salt wash buffer (20 mM Tris pH 7.5, 0.1% SDS, 1 % Triton X-100, 150 mM NaCl, 1 mM EDTA), 3 times with LiCl wash buffer (10 mM Tris pH 7.5, 1% sodium deoxycholate, 1% Triton X-100, 250 mM LiCl, 1 mM EDTA), and 1 time with TE. DNA was eluted by resuspending the beads with 125ul of elution buffer (50 mM NaHCO_3_, 1% SDS), rotating the beads for 10 minutes at room temperature followed by 3 minutes of vigorous shaking at 37 degrees. Samples were then placed on the magnet and supernatants transferred to a new tube. Elution was performed 1 additional time (250 ul total). 5 ul of 20 mg/ml Proteinase K (Roche) was added to each sample. The samples were digested and crosslinks reversed by incubating at 55 degrees for 2 hours and 65 degrees overnight. DNA was purified using the Qiagen MinElute PCR Purification Kit and ChIPseq libraries (5ug for H3K27Ac, 100ug for H3K4me3, H3K9me3 and input) were generated using the NEBNext Ultra II DNA Library Prep Kit for Illumina. Libraries were run on a NextSeq500 to generate 2×75bp reads.

#### ChIP-seq analysis

Illumina paired end reads were mapped to ERV and VL30 loci and the region 1 kb upstream from each loci using Bowtie2 (Langmead and Salzberg, 2012) with the following options: --end-to-end, --very-sensitive, and -fr. Read duplicates were removed and BAM files were generated with the Picard toolkit (Broad Institute). The BAM files were analyzed using the deepTools (Ramirez et al., 2016). Normalized bigWig files were generated using the bamCoverage tool from the BAM files using the following options: --binSize 10 --normalizeUsing RPGC -- effectiveGenomeSize 1000000 -extendReads. Normalized read counts were determined from the bigWig files using the computeMatrix tool and analyzed using R and Excel.

#### Quantitative trait locus (QTL) analysis

46 adult (8-10 week) mice from the C57BL/6N x C57BL/6J F2 intercross were genotyped by The Jackson Laboratory from ear tissue using their C57BL/6 substrain characterization panel containing 150 SNP markers spaced evenly across all chromosomes. Genotype data was examined prior to analysis and errors in genotyping (identified by expanded intermarker distances and improper linkage from an estimated recombination fraction plot) were removed. Mice were phenotyped for lymphocyte surface ERV envelope protein expression and total splenocyte ERV envelope mRNA. Single-locus QTL analysis was performed using the package R/qtl (Broman et al., 2003) using standard interval mapping with Haley Knott regression. The null distribution for the genome-wide maximum LOD score was generated by performing 10,000 permutation tests on the genotype and phenotype data. The genome-scan-adjusted p-value for each LOD peak was then calculated using an alpha of 0.05. The location of the QTL interval was estimated using the Bayes 95% credible interval and centiMorgan units were converted to base-pairs using Mouse Map Converter (Cox et al., 2009).

#### Whole genome sequencing & analysis

Genomic DNA was isolated from splenocytes using the AllPrep DNA/RNA kit or Blood & Tissue DNeasy kit (Qiagen). Library preparation and sequencing were carried out by the Yale Center for Genome Analysis. B6N and B6J samples were prepared as standard Illumina paired-end DNA libraries and used to generate 2 x 150bp reads on a HiSeq4000. All bioinformatics analyses were performed using the Ruddle High Performance Computing Cluster through the Center for Research and Computing at Yale University. Read quality was assessed by FastQC (https://www.bioinformatics.babraham.ac.uk/projects/fastqc/) and adapters were removed and reads were trimmed with Trimmomatic (Bolger et al., 2014) (LEADING:3 TRAILING:3 SLIDINGWINDOW:4:20 MINLEN:36). Reads were aligned to mm10 using BWA-mem with default settings. Alignments were sorted and indexed using SAMtools and duplicates were removed using Picard (Broad Institute). Base recalibration and variant calling were performed using GATK BaseRecalibrator and HaplotypeCaller (Van der Auwera et al., 2013) with SNP and structural variant data from the Wellcome Sanger Institute’s Mouse Genomes Project. All high-quality SNP and structural variant calls within the Bayes wide QTL interval that were not present in both B6N and B6J genomes with at least 5 total reads and an alternate allele frequency of at least 35 were analyzed with Ensembl’s Variant Effect Predictor tool and manually inspected using Integrated Genomics Viewer (Broad Institute). Additionally, chromosome 13 sequencing data was extracted using SAMtools from whole-genome sequencing data for NZB/BlNJ, NZW/LacJ, 129S5/SvEvBrd, and 129P2/OlaHsd genomes obtained from the Wellcome Sanger Institute’s Mouse Genomes Project. The copy number variant discovery programs Sequenza (Favero et al., 2015) and CNVnator (Abyzov et al., 2011) were run on all genomes for chromosome 13 using default settings.

#### 2410141K09Rik^−/−^Gm10324^−/−^ (*Snerv1/2^−/−^*) mouse generation

Two sets of guide-RNAs were designed in collaboration with the Immunobiology CRISPR Core at Yale University. B6J male mice were mated to superovulated female mice and fertilized embryos were isolated by the CRISPR core. Guide-RNAs were microinjected into the isolated embryos, which were then transferred into pseudopregnant C57BL/6J females. Seventeen pups were obtained and genotyped for CRISPR-mediated deletions.

#### Genotyping

Ear punches were obtained from mice and gDNA purified using the DNeasy Blood & Tissue Kit (Qiagen). Genotyping primers were designed to flank the sgRNA cut sites and used with TopTac Master Mix Kit (Qiagen) with 5ng of gDNA. To quantitate allele copy number, qPCR was performed as described above using 20ng of gDNA per reaction. Primer sequences are listed in Table S2. PCR-amplified products were excised, gel purified using the Zymoclean™ Gel DNA Recovery Kit (Zymo Research) and sent for sequencing at the Keck DNA Sequencing Facility at Yale University. Amplified products were also ligated into sequencing vector using the Zero Blunt^®^ TOPO^®^ PCR Cloning Kit (ThermoFisher). Competent DH5α cells were transformed, plated onto LB-agar plates containing kanamycin, and grown overnight at 37 degrees Celsius. Colonies were selected and grown overnight in 3mL of LB-kanamycin and plasmids were isolated using the QIAprep Spin Miniprep Kit (Qiagen) and sequenced.

#### TaqMan gDNA qPCR

A custom TaqMan primer-probe set (ThermoFisher) with a unique binding site within the deleted interval of interest was designed. Along with TaqMan primer-probe sets for mouse transferrin receptor and for Rybp-pseudogene, copy number assays were performed using TaqMan Genotyping Master Mix (ThermoFisher) as 20uL reactions in triplicate using 5-20ng of gDNA per reaction. qPCR data was analyzed using CopyCaller software (ThermoFisher).

#### 10x Whole Genome Sequencing

*Snerv1/2^−/−^* (C57BL/6J), *A^−/−^* (C57BL/6J), and NZB/BlNJ samples were prepared as 10x whole genome libraries and used to generated 2×150bp reads on a NovaSeq6000 by the Yale Center for Genome Analysis. The Long Ranger software pipeline (10x Genomics) was used to align reads and call structural variants.

#### Nuclear extract preparation

293T cells were transfected with 1μg of FLAG-tagged ZFP809, 2410141K09RIK (SNERV1) or GM10324 (SNERV2) expression plasmids, and 48hr later, the nuclear protein fraction was recovered by first collecting and washing cells with cold PBS using centrifugation at 1,800rpm for 10min at 4 degrees Celsius. Cell pellets were resuspended and then incubated on ice for 20min in 400uL of cold Buffer A (10mM HEPES-KOH (pH 7.9), 10mM KCl, 1.5mM MgCfe, with cOmplete Protease Inhibitor Cocktail (Roche)), after which 25uL of 10% NP-40 in Buffer A was added. The sample was vortexed at high speed for 10sec, and the homogenate centrifuged at 14,000rpm for 1min at 4 degrees Celsius. The pellet was resuspended in 1mL of Buffer A and centrifuged again at 14,000rpm for 1 min at 4 degrees Celsius. The resulting nuclear pellet was resuspended with vigorous pipetting in 100uL of cold Buffer B (20 mM HEPES-KOH (pH 7.9), 10% glycerol, 420mM NaCl, 0.2mM EDTA (pH 8.0), 1.5mM MgCl_2_, with cOmplete Protease Inhibitor Cocktail). The nuclear sample was transferred to a 1.5 ml Bioruptor Plus TPX microtube (Diagenode) and sonicated for 5 cycles of 30 seconds on/30 seconds off, on high power (Diagenode Biorupter Plus). After sonication the nuclear lysate was centrifuged for 15 minutes at 15,000rpm at 4 degrees Celsius and the clear supernatant transferred to a new tube. An equal volume of Buffer C (20mM HEPES-KOH (pH 7.9), 30% glycerol, 1.5mM MgCl_2_, 0.2mM EDTA, with cOmplete Protease Inhibitor Cocktail) was added to the extract, and the sample was stored at -80 degrees Celsius. Protein concentration was determined using the DC Protein Assay (Bio-Rad) with bovine serum albumin (Pierce) as the standard.

#### Immunoprecipitation and western blotting

100μg of nuclear extract was incubated with 2μg of anti-FLAG antibody (Sigma F1804) for 2hr at 4 degrees Celsius, and then incubated with 40μl of ProteinG-Dynabeads (Thermo Fisher Scientific) for 1hr at 4 degrees Celsius. Immunoprecipitates were washed two times with wash buffer 1 (500mM NaCl, 5mM EDTA, 20mM Tris-HCl (pH 7.5). 1% Triton-X-100) and once with wash buffer 2 (150mM NaCl, 5mM EDTA, and 20mM Tris-HCl (pH 7.5)). Samples were eluted by boiling in SDS sample buffer and subjected to electrophoresis on 10% polyacrylamide gels. PVDF blots of immunoprecipitated samples or the input fraction were probed with anti-KAP1 (Abcam ab22553), anti-FLAG, and HRP-anti-p84 (GTX70220-01) primary antibodies, and HRP-goat anti-mouse IgG (Jackson 115-035-003).

#### Recombinant protein production

2410141K09RIK (SNERV1) and GM10324 (SNERV2) open reading frames were cloned into the pGEX-6P-1 vector (Clontech) and transformed into strain C3030 (NEB). Bacteria were grown in 2xYT media and protein was expressed and batch purified by first growing bacterial cultures overnight from glyercol stocks in 2xYT medium (Sigma) with ampicillin. 250mL cultures were inoculated and grown at 25 degrees Celsius to an OD_600_ of ~0.6. Cultures were induced with 0.5mM IPTG (American Bioanalytical AB00841) and grown for 16hr at 16 degrees Celsius, and bacteria were pelleted and stored at -20 degrees Celsius. Bacteria were lysed in Lysis Buffer (50mM Tris-HCl (pH 7.4), 100mM NaCl, 0.1% Trition-X-100, 5mM DTT, with cOmplete Protease Inhibitor Cocktail) and transferred to 1.5 ml Bioruptor Plus TPX microtubes. Samples were sonicated for 7 cycles of 30 seconds on/30 seconds off, on high power. After sonication the bacterial lysates were centrifuged for 15 minutes at 14,000rpm at 4 degrees Celsius and the cleared soluble fractions were pooled in a new 15mL tube, to which 2mL of a 50% Glutathione Sepharose 4B (GE) slurry matrix was added. The sample was incubated for 1hr at 4 degrees Celsius and then washed three times with cold PBS. Protein was eluted three times with 1mL of glutathione elution buffer (50mM Tris-HCl, 10mM reduced glutathione, pH 8.0), concentrated by centrifugation (Pierce 88531), and quantitated via colorimetric protein assay (Bio-Rad). Protein fractions were run and visualized on a 10% TGX stain-free gel (Bio-Rad).

#### DNA pull-down assay

Biotinylated-ssDNA and non-labeled ssDNA were annealed via incubation at 95°C for 10 min, and then conjugated to streptavidin Dynabeads (Thermo Fisher Scientific M280) at room temperature for 1 hr in DB buffer (20mM Tris-HCl pH8.0, 2M NaCl, 0.5mM EDTA, 0.03% NP-40). 10μg of DNA-conjugated Dynabeads were incubated with 10μg of each of the FLAG-tagged recombinant proteins at RT for 30min in PB buffer (50mM Tris-HCl pH 8.0, 150mM NaCl, 10mM MgCl_2_, 0.5% NP-40, proteinase inhibitor). Beads were then washed three times with PB buffer, eluted, and subjected to immunoblotting, as described above. Oligonucleotide sequences are listed in Table S2.

#### Electrophoretic Mobility Shift Assay (EMSA)

EMSAs were performed following a published protocol (Steiner and Pfannschmidt, 2009) by first annealing AlexaFluor-488-labeled and non-labeled ssDNA to form dsDNA, as described above. Binding reactions (1x binding buffer (Thermo Fisher Scientific), 50ng/ul sonicated salmon sperm DNA (Invitrogen), 10mM MgCl_2_, 150mM NaCl, 0.5% NP-40) were mixed with 0-10ug recombinant protein and 0.9ng/uL of probe, and incubated at RT for 30min. If included, unlabeled competitor probe was added in 10-fold excess to labeled probe. Reactions were run on 6% TBE gels (Invitrogen) without loading dye. Probe migration was detected on a ChemiDoc MP Imaging System (Bio-Rad). Oligonucleotide sequences are listed in Table S2.

### Statistical Analysis

In Figure 1A-B, mean and standard deviation were plotted, with n=8 for each group. Figure 1C, Figure 1E, & Figure S1C replicates were obtained from n=2 mice for each group. Figures 1D, Figure 1F, & Figure S1D, replicates were obtained from BMDM cultures generated from pooled bone marrow from 3 B6N or B6J mice. Figure S1A replicates were obtained from separate BMDM cultures from 3 individual mice. Figure S1B replicates were obtained from individual wells of cultured mES cells. P-values in Figure 1 and Figure S1 were calculated for multiple t-tests (two-tailed) comparing B6N to B6J for each gene, corrected for the 25 independent hypotheses tested in Figure 1 and Figure S1 using the Holm-Šidák method with an alpha value of 0.05 for the entire family of comparisons. Adjusted p-values in Figure 1D-E were calculated using DESeq2. In Figure 2B, mean and standard deviation were plotted. P-values were calculated for multiple t-tests (two-tailed) comparing NEERV to VL30 for each histone mark across each region, corrected for the 9 independent hypotheses tested using the Holm-Šidák method with an alpha value of 0.05 for the entire family of comparisons. P-values in Figure S2C were calculated using one-way ANOVA with an alpha value of 0.05, 45 degrees of freedom, and F-values of 1.35 (Xmv), 0.207 (Pmv), 0.274 (B-cell MFI), 0.414 (CD4 T-cell MFI), and 0.067 (CD8 T-cell MFI). In Figure 3A, mean and standard deviation were plotted, with n=4 or n=8 for each group. P-values in Figure 3A were calculated using one-way ANOVA with Šidák’s multiple comparisons test, an F-value of 16.49, 17 degrees of freedom, and an alpha value of 0.05. Genome-wide P-values for the QTL analysis were calculated by performing 10,000 permutation tests to obtain a genome-wide distribution for the null hypothesis. LOD thresholds for an alpha value of 0.05 were calculated as 3.72 (Xmv), 3.95 (Pmv), 3.90 (B-cell MFI), 3.85 (CD4 T-cell MFI), and 3.68 (CD8 T-cell MFI). In Figure 4C, mean and standard deviation were plotted, with n=10 (B6N), n=10 (B6J), n=16 *(241Rik^−/−^Gm10324^−/−^*), n=18 (*241Rik^-/+^Gm10324^-/+^*), n=9 (A^−/−^), n=12 (A^+/-^), n=9 (B^−/−^), n=5 (B^+/-^), n=12 (C^−/−^), n=7 (C^+/-^), n=17 (B6J littermates). In Figure 4E mean and standard deviation were plotted with n=7 (B6J, *241Rik^−/−^Gm10324^−/−^)* and n=8 (B6N). P-values for Figures 4C & 4E were calculated for multiple t-tests comparing all genotypes to the B6J WT littermate value for each gene (Figure 4C) or comparing the *241Rik^−/−^Gm10324^−/−^* value to that of B6J (Figure 4E), corrected for the 33 independent hypotheses tested using the Holm-Sidák method with an alpha value of 0.05 for the entire family of comparisons. Adjusted p-values in Figure 4D & Figure S4D were calculated using DESeq2. In Figure 6A-F, mean and standard deviation were plotted. In Figure 6A, n=14 *(Snerv1/2^−/−^NZB*), n=19 (B6JxNZB). In Figure 6B-C, n=19 (Snerv1/2^−/−^xNZB) and n=23 (B6JxNZB). In Figure 6D-F, n=16 for each group. P-values were calculated for multiple t-tests comparing the Snerv^/-^-based F1 value to the B6J-based F1 value for each gene, corrected for the 20 independent hypotheses tested using the Holm-Šidák method with an alpha value of 0.05 for the entire family of comparisons. Adjusted p-values in Figure S6B were calculated using DESeq2. Data was analyzed using GraphPad Prism 7.

### Data and Software Availability

The BioProject accession number for sequencing data generated in this study is PRJNA498070. The Mendeley dataset is available at https://data.mendeley.com/datasets/p3bpmhtwwp/draft?a=5ff9a586-cd84-4114-b88f-77bcb7bc84b6. The mm10 locations of proviral ERV loci are listed in Table S3. The parsing and mapping algorithms used to analyze mouse proviral ERV expression in RNA-sequencing data can be found as Perl scripts in the Supplementary Information as Data S1.pl and Data S2.pl.

## Supplemental Information

### Supplemental Files

Figures S1–S7, Table S1–S3 (separate PDF)

Data S1.pl BWA BAM parsing script

Data S2.pl ERV mapping script

### Declaration of interests

The authors declare no competing interests.

---

**Figure S1.**
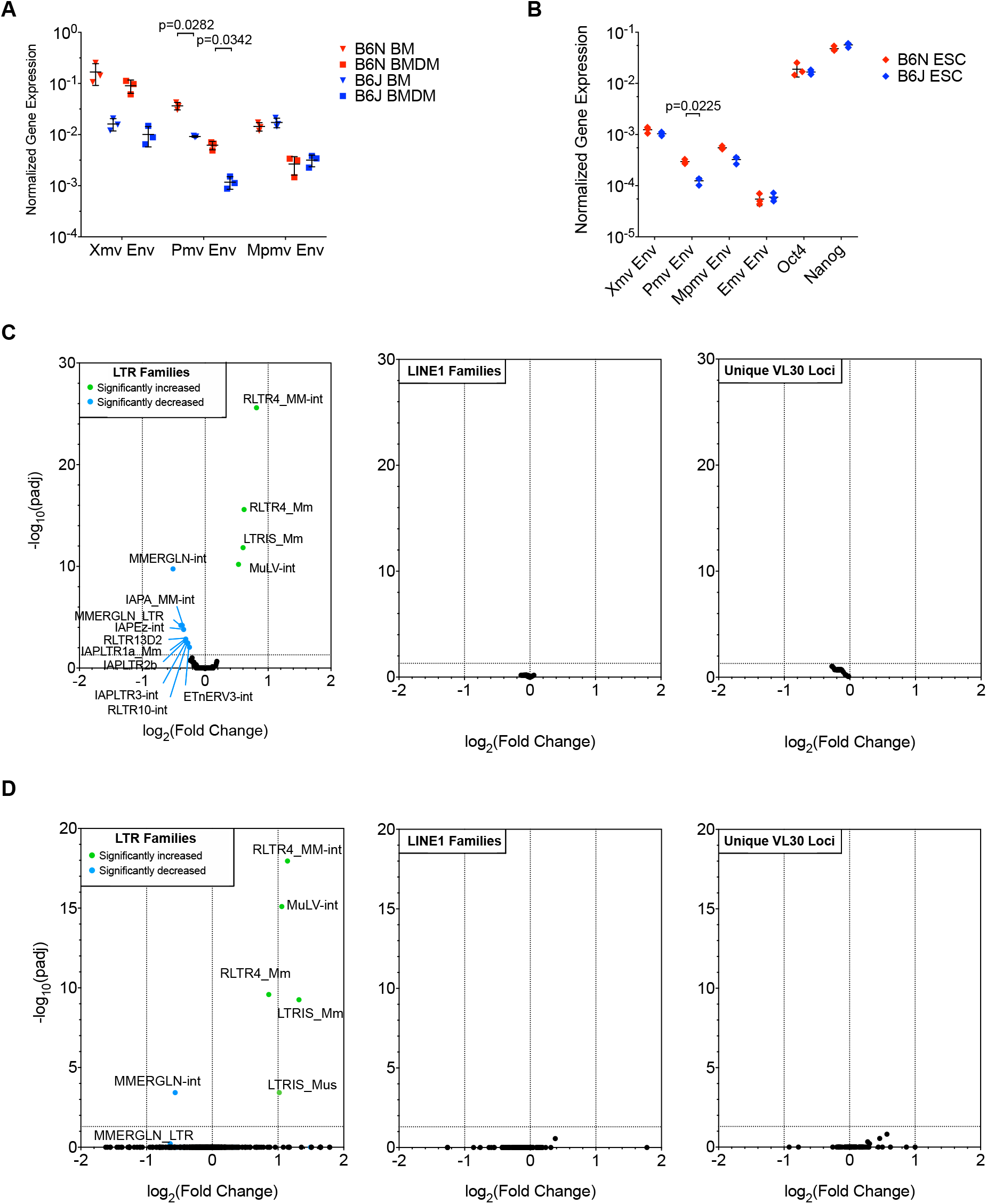
Increases in endogenous retrovirus transcription in C57BL/6N, but not C57BL/6J, mice do not depend on cell type or differentiation state, Related to Figure 1. **(A)** RT-qPCR of RNA from total bone marrow (BM, n=3) and bone marrow-derived macrophages (BMDM), n=3 from B6N and B6J mice. **(B)** RT-qPCR of RNA from B6N and B6J embryonic stem cells cultured in triplicate off of B6J feeders for 48hr. Oct4 and Nanog serve as pluripotency markers. **(C-D)** Volcano plot of all 680 LTR families or 132 L1 families listed in RepBase, or unique VL30 loci from 2×150bp mRNA sequencing of B6N and B6J naïve CD4^+^ T cells (C) and bone marrow-derived macrophages (D). Figure S1A-B adjusted p-values were calculated for multiple t-tests comparing B6N to B6J for each gene, corrected for the 25 independent hypotheses tested in Figure 1 and Figure S1 using the Holm-Šidák method with an alpha value of 0.05 for the entire family of comparisons. Adjusted p-values from DESeq2 are reported for RNA-sequencing results.

**Figure S2.**
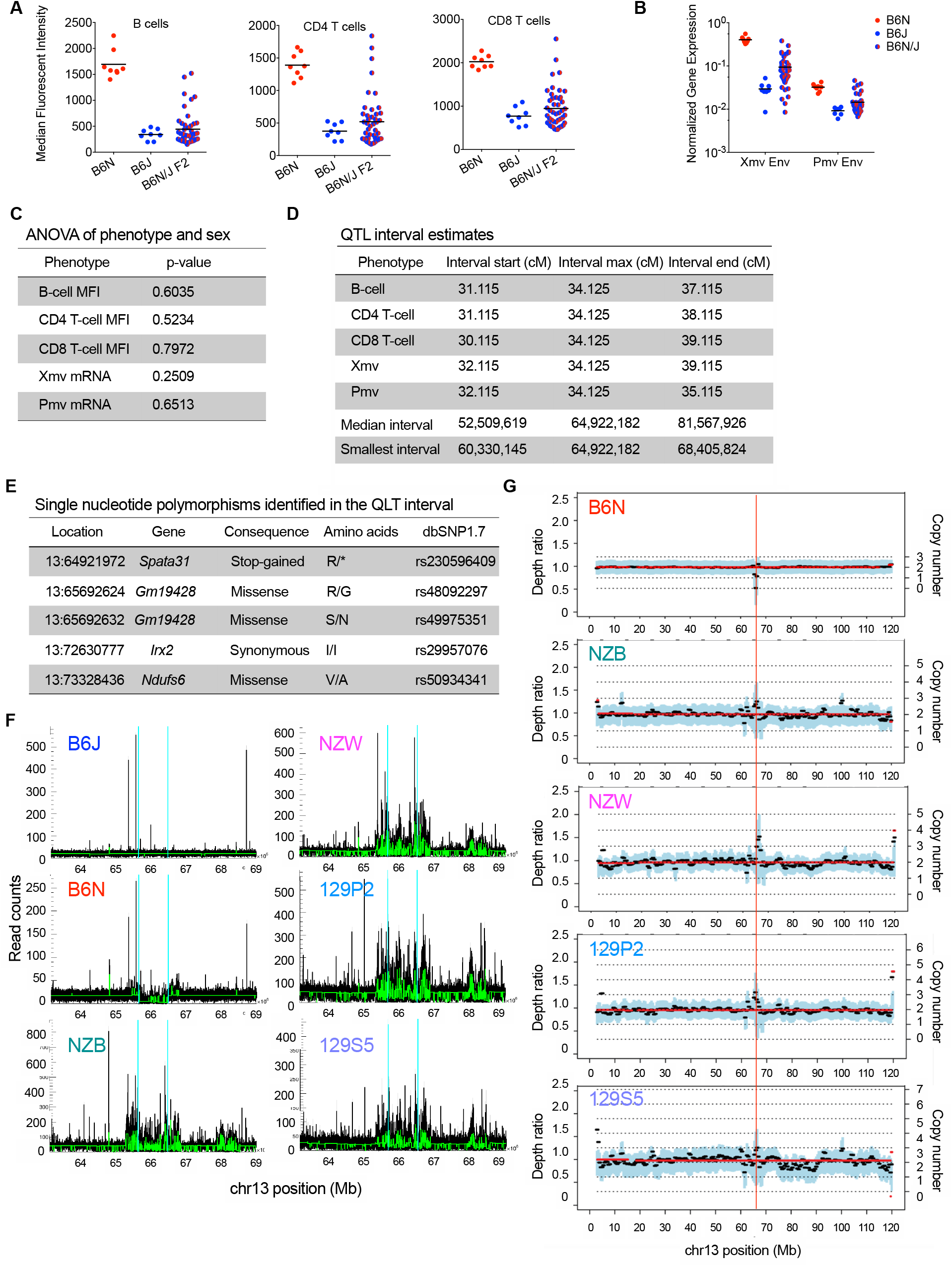
The C57BL/6NJ F2 intercross data gives rise to consistent chromosome 13 QTL interval estimates, and *in silico* analyses indicate that a large deletion is present within this QTL interval in the C57BL/6N genome, Related to Figure 3. **(A)** Calculated MFI of ERV envelope protein expression detected via FACS on the surface of peripheral blood B cells, CD4^+^ T, and CD8^+^ T lymphocytes or **(B)** RT-qPCR of RNA from total splenocytes from B6N (n=8), B6J (n=8), and C57BL/6NJ F2 (n=46) mice. **(C)** One-way analysis of variance results value of each QTL phenotype by gender. **(D)** QTL interval estimates for each phenotype using the Bayes 95% credible interval. **(E)** Location and consequence of coding single nucleotide polymorphisms identified within the QTL interval from B6N whole genome sequencing data. **(F)** CNVnator determines copy number variants based upon read-depth analysis within a sample, with average read count drawn in green. CNVnator identifies a single significant structural variant within the QTL interval in the B6N genome, a ~840,000bp loss in read depth (demarcated by vertical blue lines). CNVnator analysis of this region in NZB, NZW, 129S5, and 129P2 genomes is also shown. **(G)** Sequenza analysis of chromosome 13 was performed on B6N, NZB, NZW, 129S5, and 129P2 whole genome sequencing data. Sequencing depth ratio and calculated copy number are shown on the left and right axes, respectively. The position of the loss of copy number that was identified in the B6N genome is demarcated by a vertical red line. P-values in Figure S2C were calculated using one-way ANOVA with an alpha value of 0.05 and 45 degrees of freedom.

**Figure S3.**
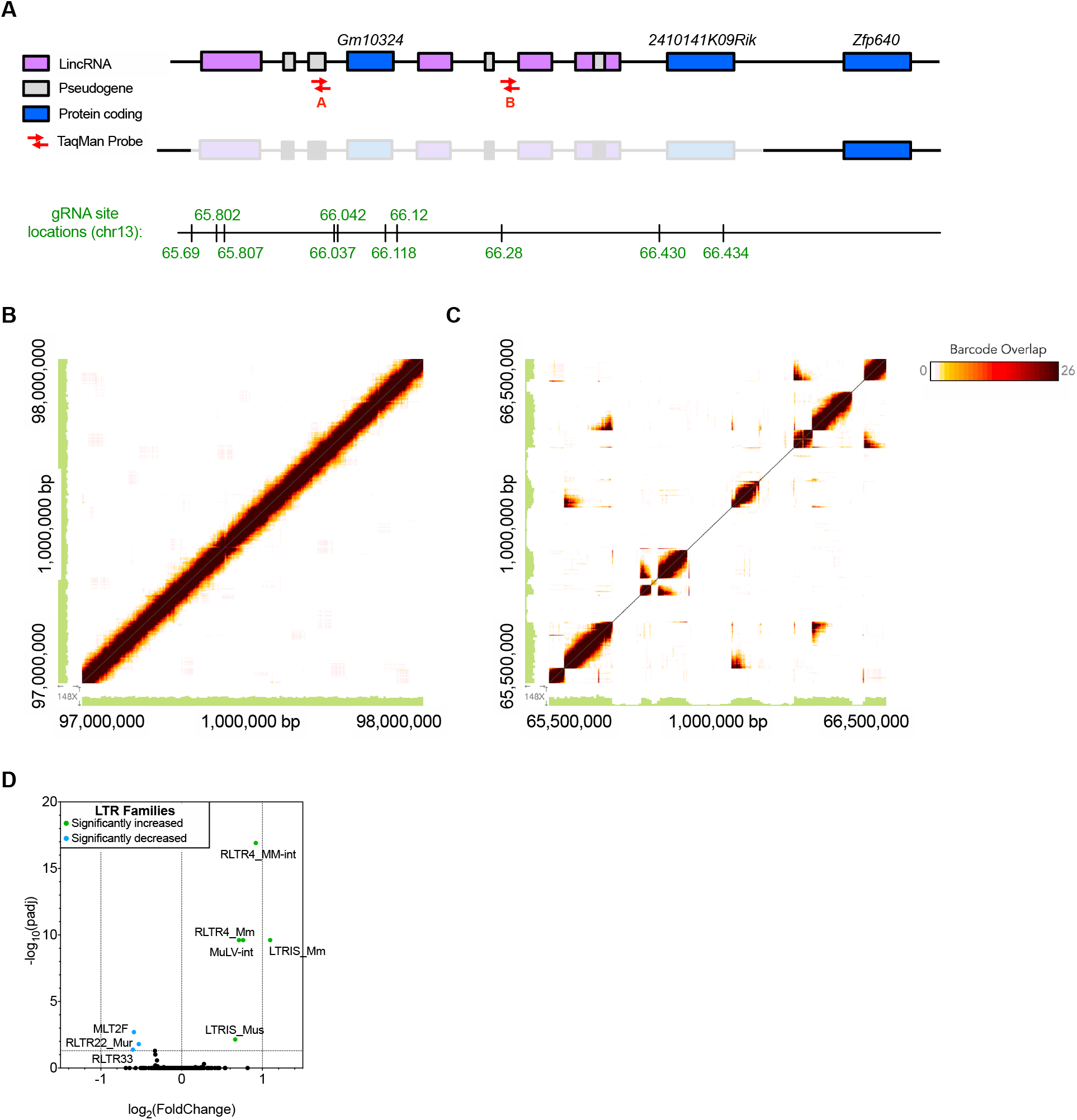
Deletion of 2 KRAB-ZFP genes from the B6J genome leads to selective increases in transcription of NEERV LTR families, Related to Figure 4. **(A)** Schematic of the locations on chromosome 13 that were targeted by CRISPR guide RNAs. **(B)** Representative Loupe software (10x Genomics) output from *241Rik^−/−^Gm10324^−/−^* 10x WGS visualizing a 1,000,000bp region from chromosome 1. The genomic coordinates of the region being viewed are listed at either end of the left and bottom axes. Adjacent DNA sequences share barcodes, resulting in high barcode overlap signal across the diagonal of the square plot. Distant DNA sequences do not share barcodes. Sequencing coverage is shown in the green histogram at the left and bottom edges of the plot. **(C)** Loupe software output from *241Rik^−/−^ Gm10324^−/−^* 10x WGS, showing the chromosome 13 region targeted by CRISPR guide RNAs. Deletions in the region are captured as gaps on the diagonal, with high barcode overlap between distant DNA regions. **(D)** Volcano plot of all 420 LTR families listed in RepBase from 2×75bp mRNA sequencing of B6J and *241Rik^−/−^Gm10324^−/−^* (B6J) CD4^+^ T cells. Adjusted p-values were calculated using DESeq2.

**Figure S4.**
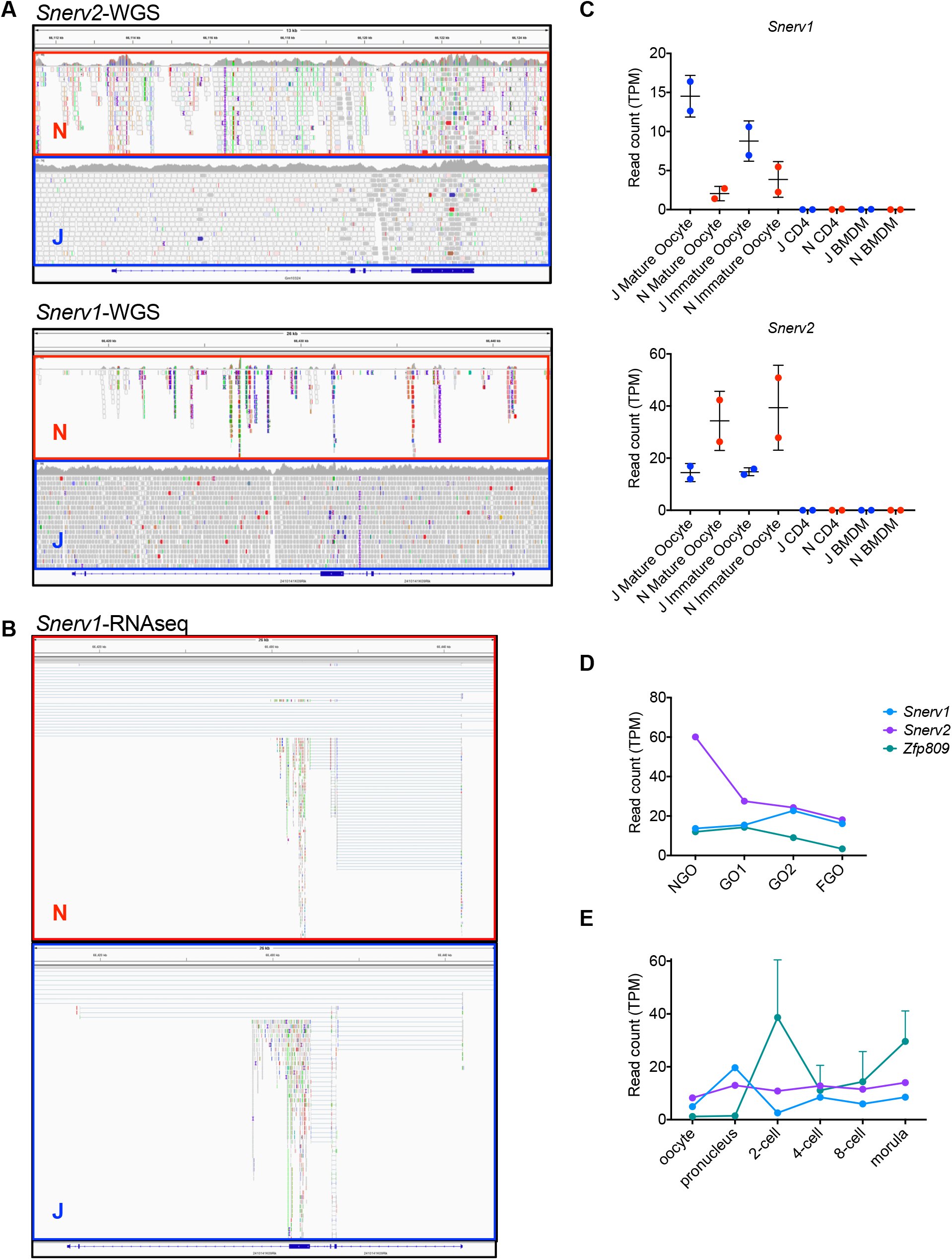
Low-confidence mapping to *Snerv1* and *Snerv2* reveals their expression in oocytes and early embryogenesis, Related to Figure 4. **(A)** Integrative Genomics Viewer (IGV; Broad Institute) output showing B6N (red) and B6J (blue) WGS reads that map to *Snerv1 (2410141K09Rik)* and *Snerv2 (Gm10324)* genes. High confidence alignments are colored gray, while low confidence alignments (e.g. more than one possible location for the alignment) are white. Colored alignments indicate mate pairs that map to a distant region of the same chromosome or to a different chromosome. Sequencing coverage is shown in the gray histogram that runs along the top of each panel. **(B)** IGV output showing B6N (red) and B6J (blue) RNA sequencing reads from B6N (red) and B6J (blue) mature oocytes that map to *Snerv1* and *Snerv2* mRNA. Reads are colored as described above, while reads with mapping locations that bypass the gene are shown as horizontal lines. **(C)** Read counts in transcripts per kilobase million (TPM) for *Snerv1* and *Snerv2* mRNA from B6N (red) and B6J (blue) mature oocytes, immature oocytes, CD4 T cells, and BMDM. **(D)** Read counts in TPM for *Snerv1* and *Snerv2* mRNA from publicly available (Veselovska et al., 2015; Xue et al., 2013) sequencing data of B6J non-growing oocytes (NGO), growing oocytes (GO1 & GO2), and fully-grown oocytes (FGO). ZFP809 values are shown for reference. **(E)** Read counts in TPM for *Snerv1* and *Snerv2* mRNA from publicly available single-cell sequencing data of B6J oocytes (2 cells sequenced), pronucleus, 2-cell, 4-cell, and 8-cell embryos (3 cells each sequenced). *Zfp809* values are shown for reference.

**Figure S5.**
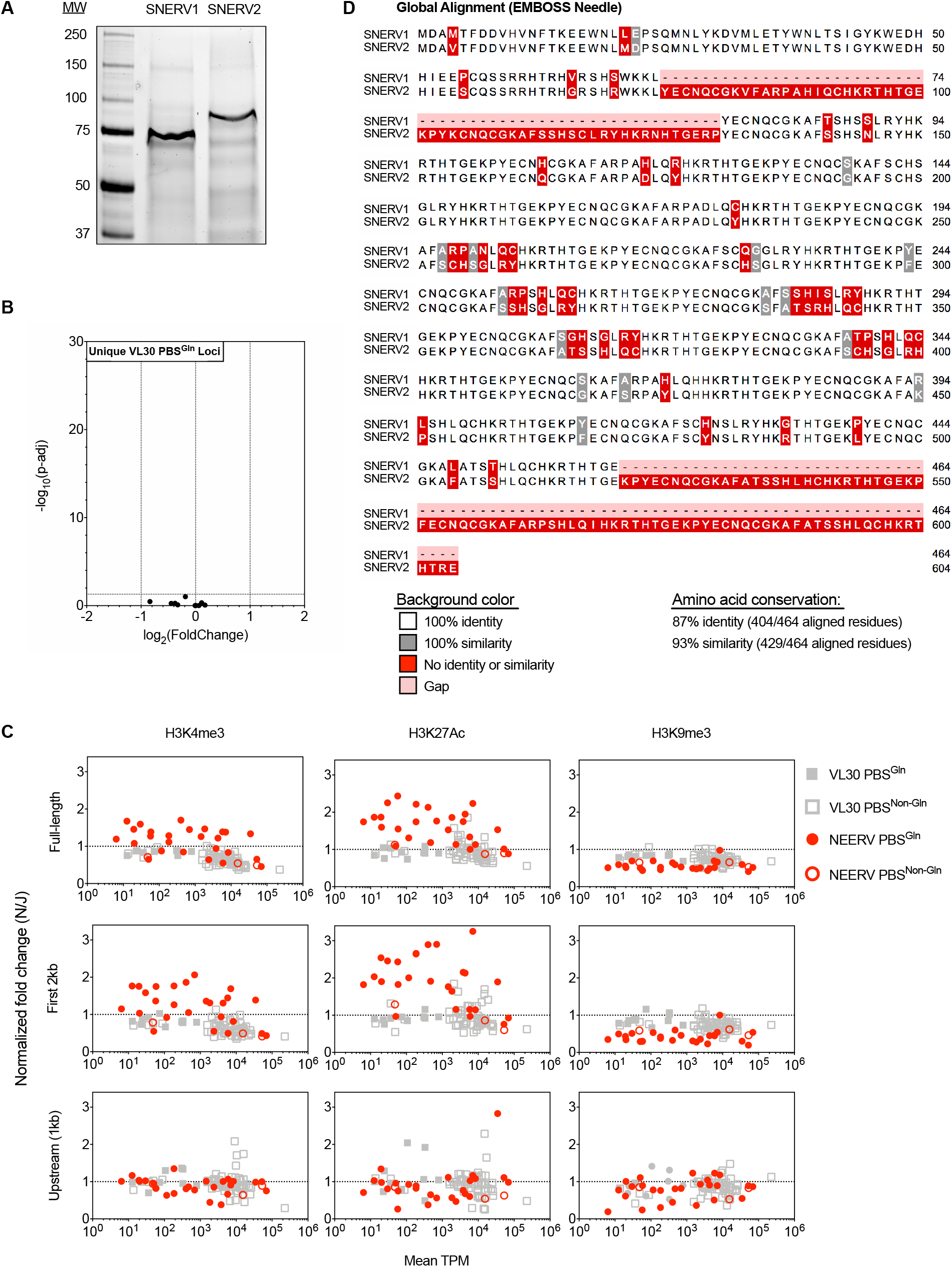
Presence of PBS^Gln^ is not sufficient for transcriptional repression, enrichment of activating histone modifications, or depletion of repressive histone modifications by SNERV1, which shares high amino acid sequence homology with SNERV2, Related to Figure 5. **(A)** Protein gel of purified recombinant GST-FLAG-SNERV1 and GST-FLAG-SNERV2 used in DNA pulldown and EMSA experiments. **(B)** Volcano plot of unique VL30 PBS^Gln^ loci from 2×150bp mRNA sequencing of B6J and *Snerv1/2^−/−^* CD4^+^ T cells. **(C)**. Plot of normalized fold change at PBS^Gln^ and PBS^Non-Gln^ NEERV and VL30 loci for each listed histone modification versus the mean expression level in B6N and B6J BMDMs in transcripts per million (TPM), as in Figure 2. **(D)** Amino acid sequences of SNERV1 and SNERV2 proteins were aligned using EMBOSS Needle Global Alignment software. Amino acid conservation for the 464 residues that align is listed. Adjusted p-values were calculated using DESeq2.

**Figure S6.**
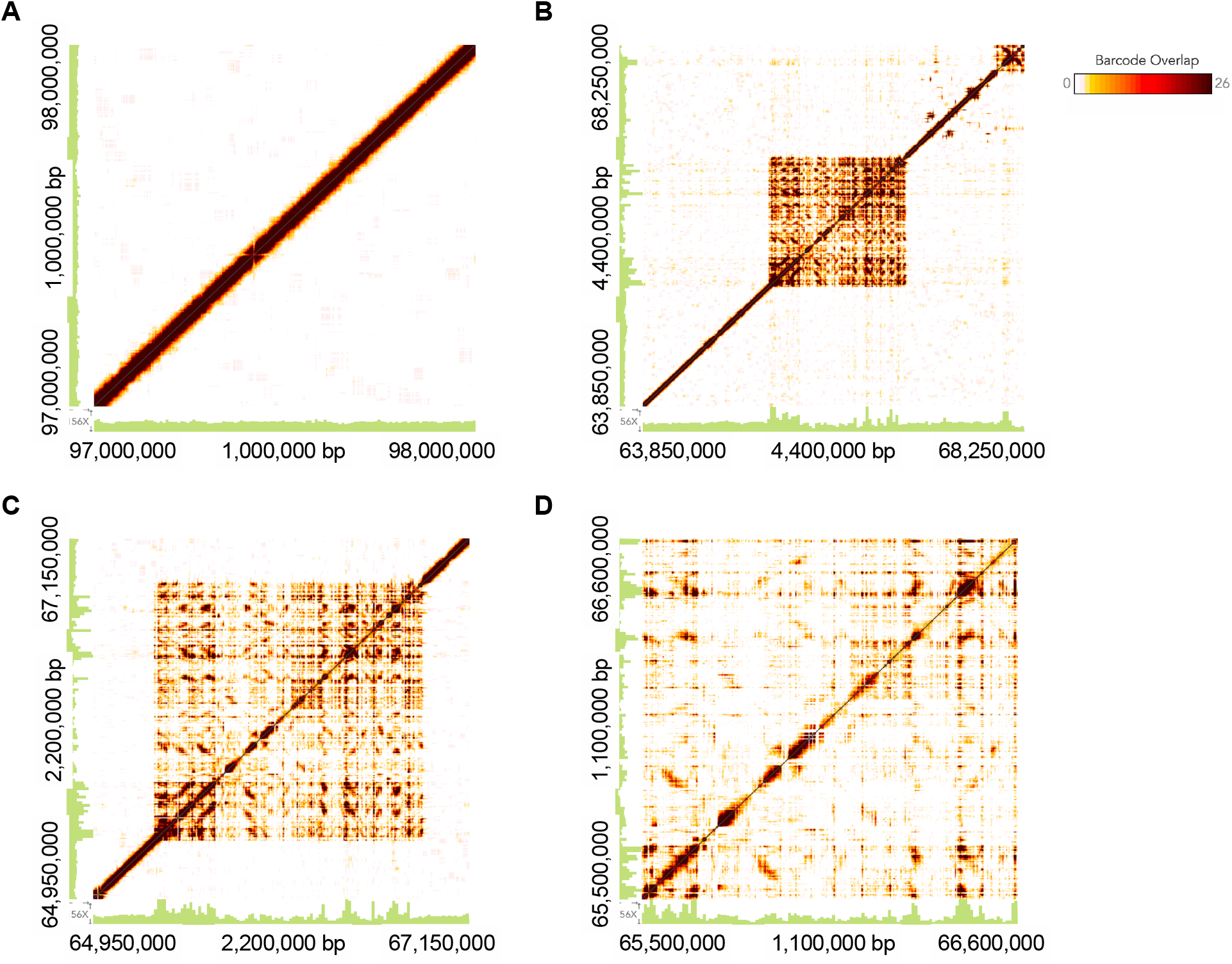
10x WGS of chromosome 13 in the NZB genome does not elucidate the structure of the *Snerv1/2-containing* region, Related to Figure 6. **(A)** Representative Loupe software (10x Genomics) output from NZB 10x WGS visualizing a 1,0, 000bp region from chromosome 1. The genomic coordinates of the region being viewed are listed at either end of the left and bottom axes. Adjacent DNA sequences share barcodes, resulting in high barcode overlap signal across the diagonal of the square plot. Distant DNA sequences do not share barcodes. Sequencing coverage is shown in the green histogram at the left and bottom edges of the plot. **(B-D)** Loupe software output from NZB 10x WGS visualizing the region on chromosome 13 containing the *Snerv1* and *Snerv2* genes.

**Figure S7.**
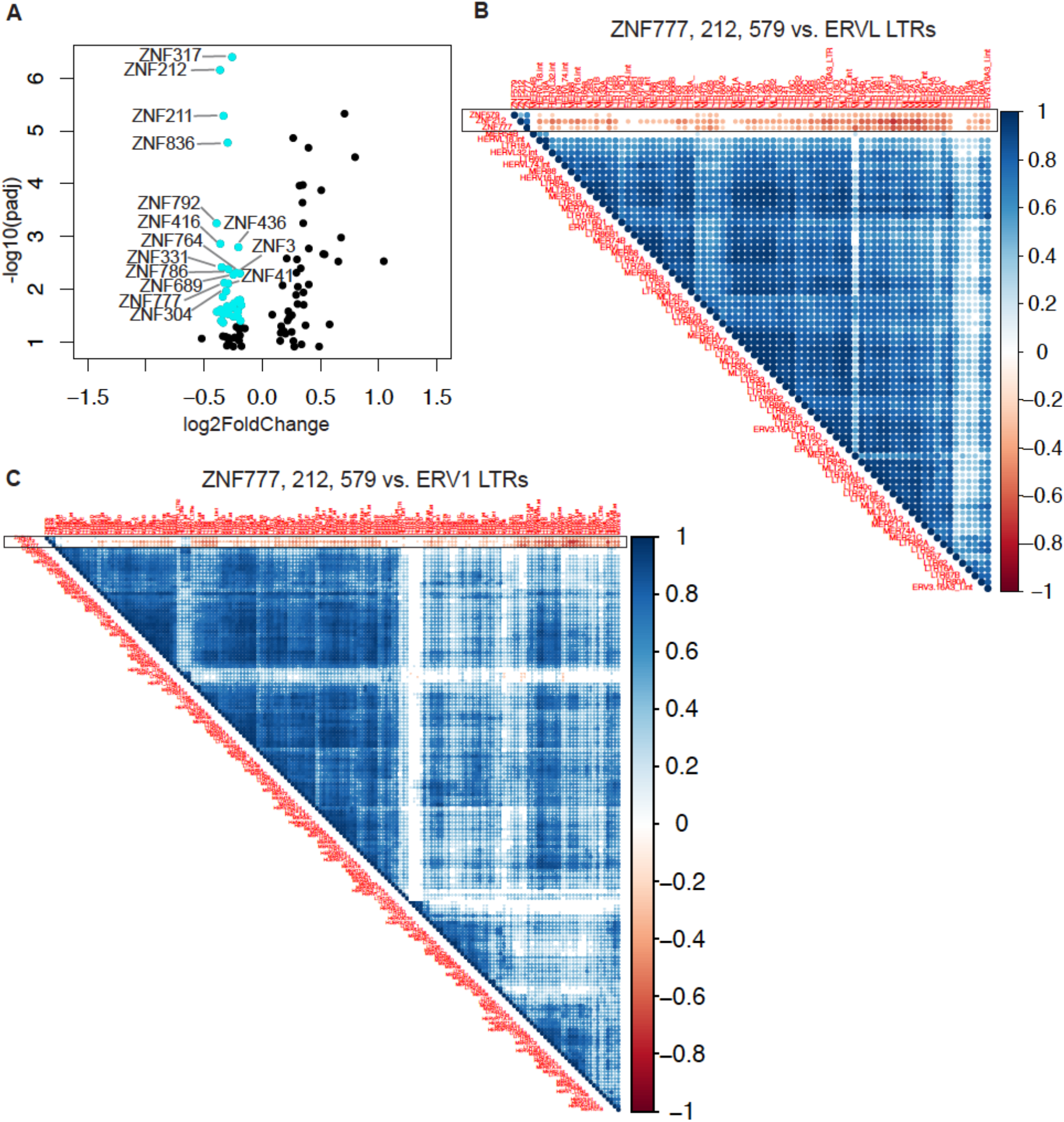
Human KRAB-ZFP ZNF212, ZNF579, and ZNF777 transcripts are significantly decreased and anticorrelate with increased HERV transcription in SLE patients, Related to Figure 7. RNA sequencing data obtained from whole blood of SLE patients (n=99) and healthy controls (n=18) were used to perform RepEnrich and DESeq2 analyses to quantify expression of LTR elements and cellular genes, respectively. **(A)** Volcano plot of KRAB-ZFP with significantly decreased expression in the blood of SLE patients versus healthy controls. LTR subfamilies indicated in aqua are log_2_(Fold Change) > 1 and padj < 0.05 in SLE patients versus healthy controls. **(B-C)** Spearman correlation was calculated between ERVL **(B)** and ERV1 **(C)** LTR subfamilies and ZNF579, ZNF212, and ZNF777 among SLE patients. The three rows corresponding to ZNF579, ZNF212, and ZNF777 are demarcated by a black box. The correlation plot represents spearman r values and displays only correlations that were p < 0.05. Blank indicates not significant.

**Table S1.**
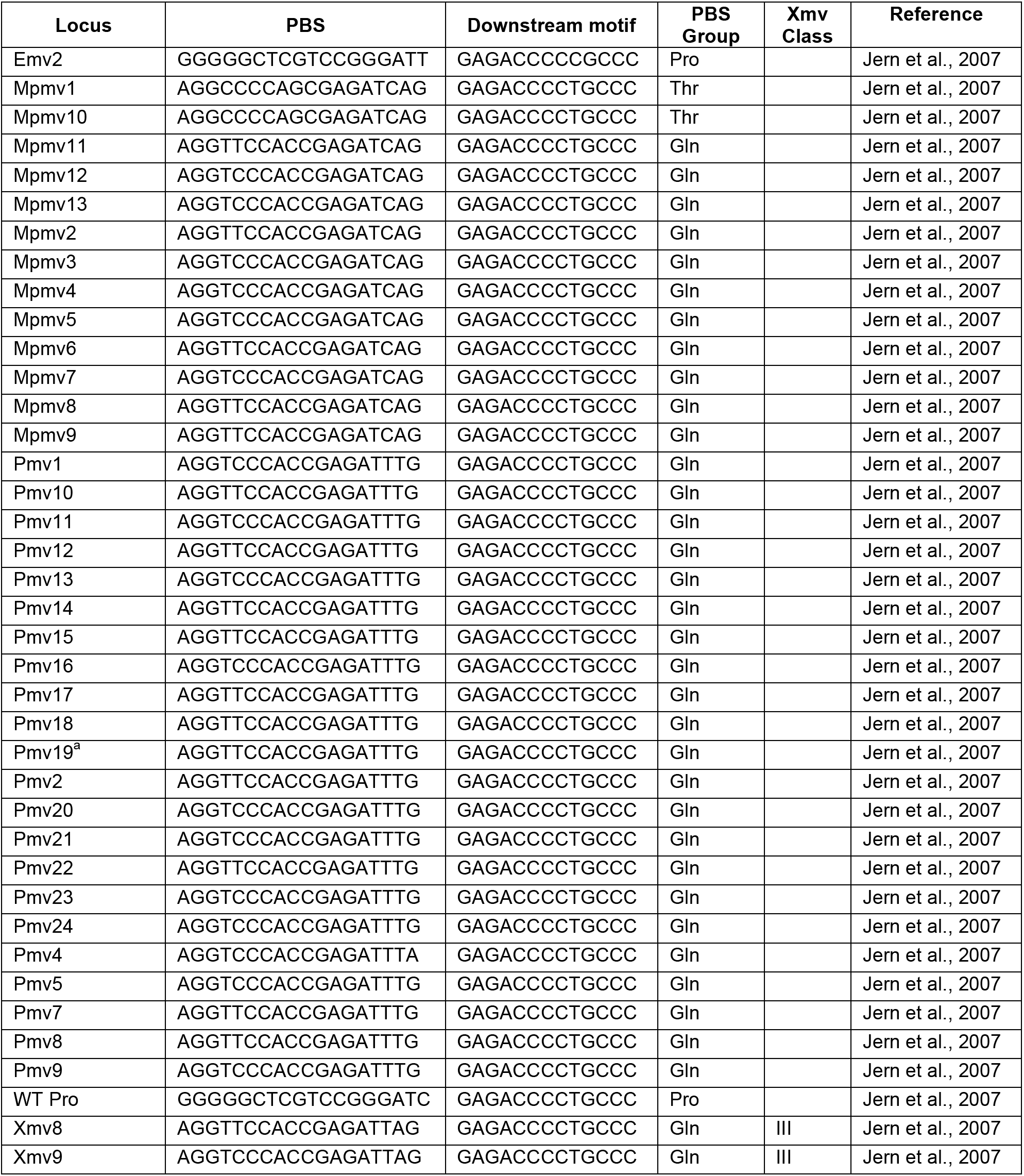
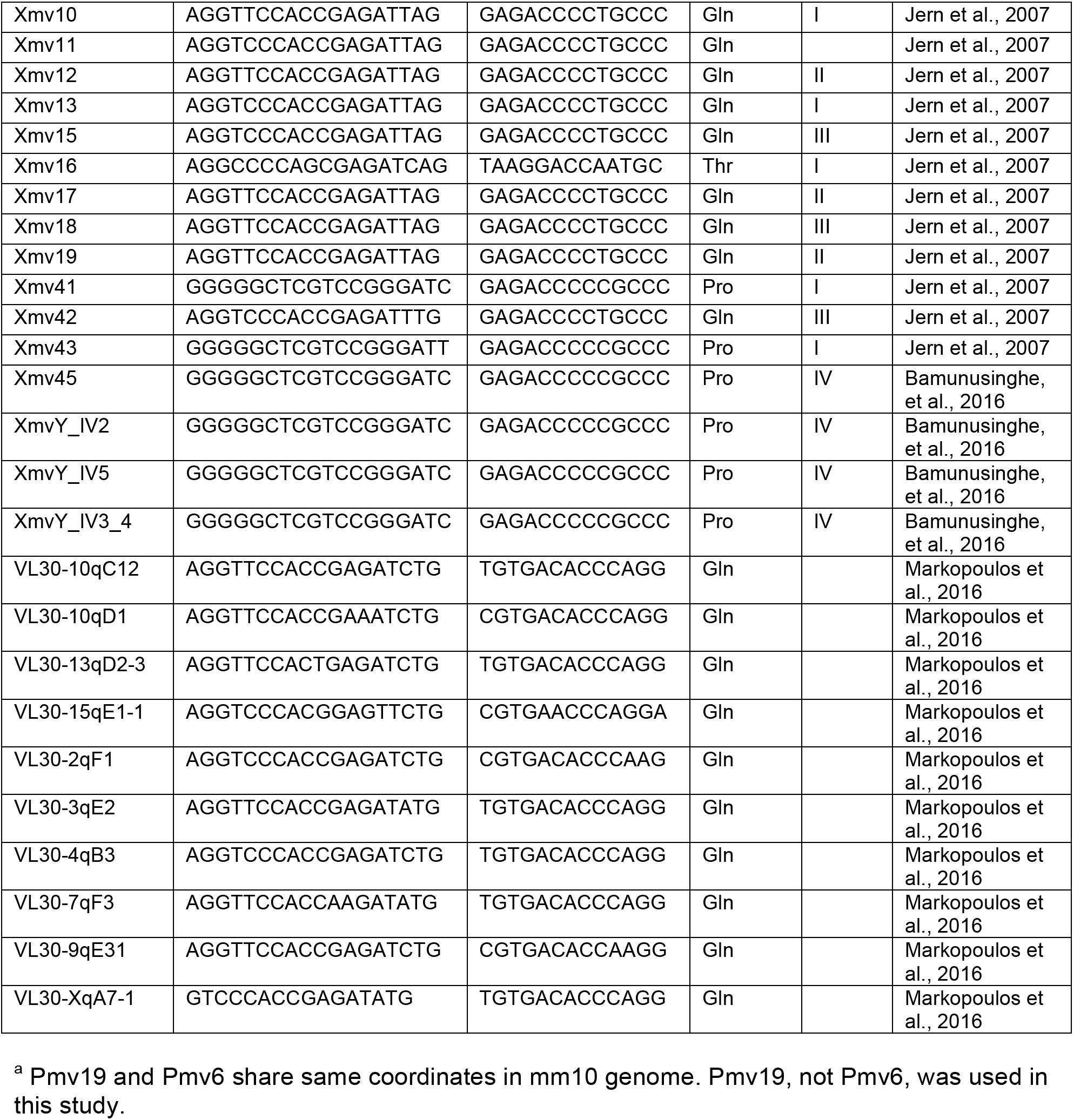
Primer binding site sequences, downstream motifs, and Xmv classes (when applicable) of unique C57BL/6 proviral endogenous retroviruses and select VL30 elements, Related to Figure 5, Figure 6, and STAR Methods

**Table S2.**
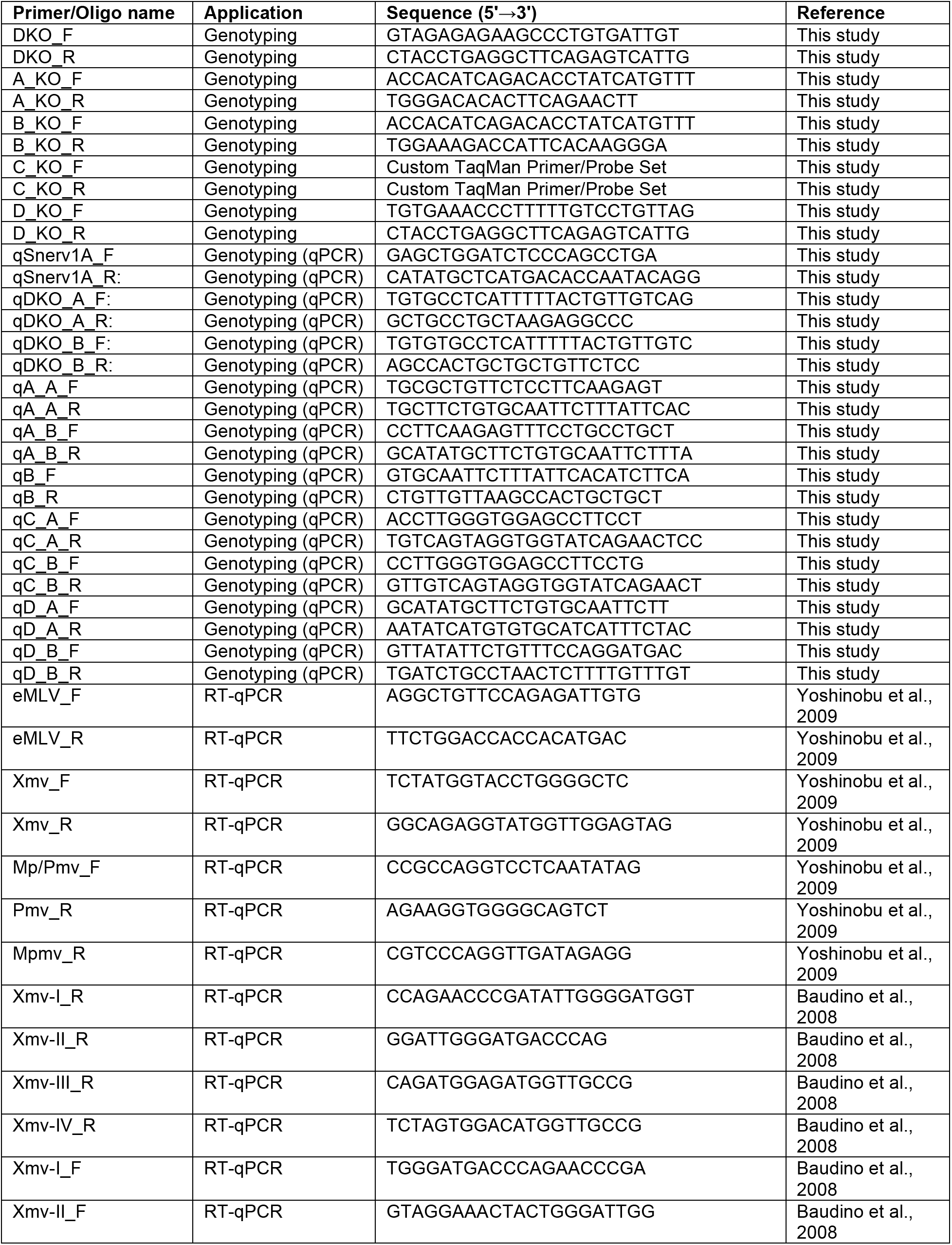
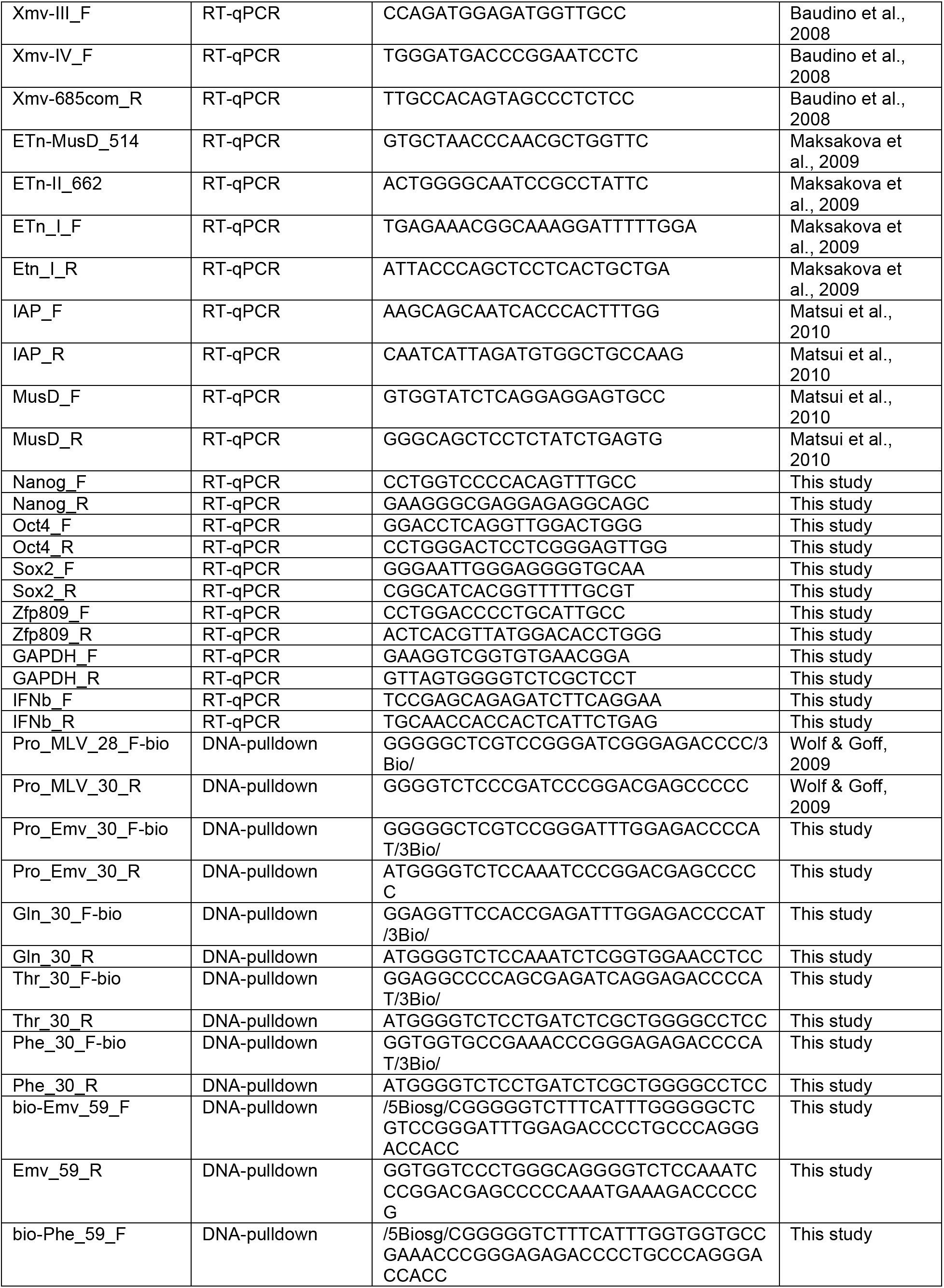
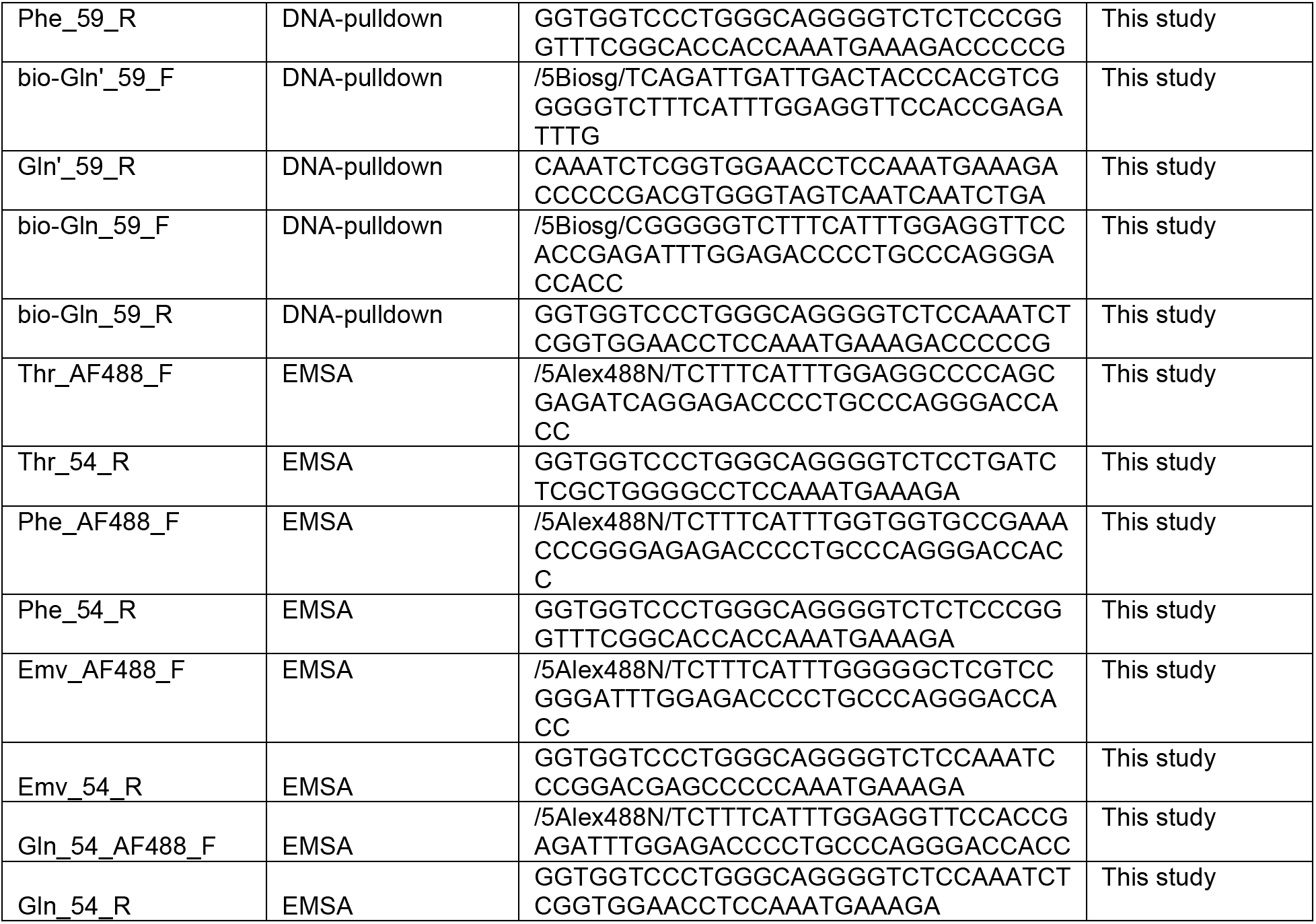
List of primers and oligos used in study, Related to STAR Methods

**Table S3.** Proviral ERV genome coordinates (mm10), Related to STAR Methods

| Locus | Chromosome | Start | End | Strand |
| --- | --- | --- | --- | --- |
| Mpmv6 | chr1 | 131642566 | 131651620 | + |
| Xmv43 | chr1 | 170941520 | 170950177 | + |
| Xmv41 | chr1 | 171481145 | 171489815 | + |
| Pmv24 | chr1 | 182257119 | 182266100 | + |
| Pmv21 | chr1 | 191933612 | 191942593 | + |
| Mpmv3 | chr2 | 16020494 | 16029536 | - |
| Pmv7 | chr2 | 57214411 | 57223392 | - |
| Xmv10 | chr2 | 156356555 | 156363605 | - |
| Mpmv10 | chr3 | 67096077 | 67103794 | + |
| Mpmv9 | chr3 | 152323139 | 152332182 | + |
| Pmv23 | chr4 | 101863156 | 101872138 | - |
| Pmv19 | chr4 | 108146033 | 108155016 | - |
| Pmv6 | chr4 | 108146033 | 108155016 | - |
| Xmv8 | chr4 | 146503082 | 146511841 | + |
| Xmv9 | chr4 | 147218827 | 147227549 | + |
| Xmv17 | chr5 | 24215307 | 24222153 | + |
| Mpmv13 | chr5 | 25230698 | 25239670 | + |
| Mpmv7 | chr5 | 43207940 | 43216981 | + |
| Pmv5 | chr5 | 44134359 | 44143341 | + |
| Pmv11 | chr5 | 76930515 | 76939497 | + |
| Pmv12 | chr5 | 144669838 | 144678819 | + |
| Pmv4 | chr7 | 7069910 | 7078892 | - |
| Pmv18 | chr7 | 29615206 | 29623905 | + |
| Pmv15 | chr7 | 30688639 | 30697620 | + |
| Mpmv1 | chr7 | 64126561 | 64135602 | + |
| Xmv12 | chr8 | 43311729 | 43320463 | - |
| Pmv10 | chr8 | 119889564 | 119897040 | - |
| Emv2 | chr8 | 123425506 | 123434150 | - |
| Xmv16 | chr9 | 41905484 | 41912268 | - |
| Xmv15 | chr9 | 62439173 | 62447932 | + |
| Pmv1 | chrX | 15474047 | 15483028 | + |
| Mpmv5 | chr10 | 5717714 | 5726753 | + |
| Mpmv12 | chr10 | 22734277 | 22743322 | + |
| Pmv13 | chr10 | 41436647 | 41445628 | + |
| Pmv8 | chr10 | 50419548 | 50427575 | - |
| Pmv2 | chr11 | 6749421 | 6758391 | - |
| Pmv22 | chr11 | 8920297 | 8929277 | + |
| Mpmv2 | chr11 | 76548807 | 76557852 | + |
| Mpmv4 | chr11 | 86882315 | 86891356 | + |
| Xmv42 | chr11 | 88890179 | 88896966 | - |
| Mpmv8 | chr11 | 103083662 | 103092704 | + |
| Mpmv11 | chr12 | 69546805 | 69555846 | + |
| Xmv13 | chr13 | 67884887 | 67893574 | - |
| Pmv9 | chr13 | 98986274 | 98995255 | - |
| Xmv19 | chr14 | 54530364 | 54538188 | + |
| Pmv17 | chr15 | 76559816 | 76567002 | - |
| Pmv14 | chr16 | 76135042 | 76144024 | - |
| Pmv16 | chr16 | 93697456 | 93706437 | - |
| Pmv20 | chr18 | 82693550 | 82702430 | + |
| Xmv18 | chr19 | 60925306 | 60934033 | - |

